# Molecular anatomy of the entorhinal cortex

**DOI:** 10.64898/2026.10.13.758661

**Authors:** Ana Morello-Megias, Bhanu Shankar Dhulipalla, Han Ali Kahvecioglu, Jerald Shin, Saiyan Joseph, Devika Aggarwal, Jennifer I. Luebke, Douglas L. Rosene, Christina Dimovasili, Jean-Pierre Roussarie

## Abstract

The entorhinal cortex (EC) is critical for episodic memory formation and is the epicenter of neurofibrillary tangle formation in Alzheimer’s disease. Its cytoarchitectonic and functional heterogeneity has been recognized for decades, but the molecular underpinnings of this heterogeneity are still relatively uncharted. While many features of the region appear to be conserved between rodents and primates, very little is known about the conservation of the molecular identity of EC neurons. Here we perform single-nucleus RNAseq (snRNAseq) of the mouse EC and use in situ hybridization, as well as publicly available transcriptomics data, to enhance the understanding of the heterogeneity of neuron types present in the area. While the well-known dichotomy between lateral and medial EC is evident in superficial layers of EC, we highlight that the medioventral region of EC forms a different population from the caudomedial section. We identify molecular correlates for the previously described parcellation of lateral EC, and show that, at least in layer IIa, subterritories of EC are not separated by abrupt borders but rather form smooth gradients. Lastly, we show that both lateral vs. medial ECIIa markers, and markers that reveal finer grain heterogeneity within ECIIa have conserved patterns of expression in macaque, highlighting the high degree of homology between the two species. Overall this molecular atlas of the mouse EC with some comparative elements is a resource for anatomically-informed molecular studies of the EC and lays the groundwork for mechanistic studies of selective neuronal vulnerability of the region in the context of AD.

## Introduction

The entorhinal cortex (EC) is a brain region at the interface between neocortex and hippocampus. It is as central for key cognitive processes like episodic memory (Scoville and Milner, 2000; Lipton and Eichenbaum, 2008) as it is in Alzheimer’s disease (AD) pathogenesis (Arnold et al., 1991; Braak and Braak, 1991, 1995; Fukutani et al., 1995, 2000; Gómez-Isla et al., 1996), where it is the prime site for deposition of tau proteinaceous aggregates in prodromal disease stages. For these reasons, a very abundant literature describes EC cytoarchitectonic features (Amaral et al., 1987; Insausti et al., 1995, 1997; Groen, 2001), its afferent and efferent connectivity to cortex and hippocampus (Insausti et al., 1997; Burwell and Amaral, 1998; Groen et al., 2003; Canto et al., 2008; Suh et al., 2011; Witter et al., 2017; Ohara et al., 2023), intrinsic electrophysiological properties of its cells (Alonso and Klink, 1993; Tahvildari and Alonso, 2005; Giocomo and Hasselmo, 2008, 2009; Canto and Witter, 2012a, 2012b), as well as network-level characteristics (Stensola et al., 2012; Fernández-Ruiz et al., 2021). Much less is known, though, about the molecular makeup of its neurons and about its transcriptional heterogeneity.

The laminar architecture of EC makes it a periallocortical region, in between the three layers of primitive allocortical regions like the hippocampus, and the 6 layers of neocortex. It has 6 layers like neocortex but its layer IV is relatively devoid of cells and called the *lamina dissecans* (Insausti et al., 2017). The EC forms the majority of the afferent inputs into the hippocampus via two different pathways: a) the perforant pathway, the projection from principal neurons of layer IIa (Hjorth-Simonsen and Jeune, 1972) (we use the nomenclature defined and used by Witter in (Witter et al., 2017; Kobro-Flatmoen and Witter, 2019; Ohara et al., 2019)) synapsing onto dendrites of dentate gyrus granule cells and CA3 pyramidal cells; and b) the temporoammonic pathway, the projection of principal neurons from layer III terminating onto dendrites of CA1 pyramidal cells in *stratum lacunosum moleculare* (Steward, 1976). The EC is also one of the main recipients of outputs of the hippocampus with CA1 pyramidal cells and subiculum neurons innervating EC neurons in layer V.

Along all these dimensions, substantial levels of heterogeneity are found across the region. The main subdivision commonly made in the mouse EC is between lateral and medial areas (LEC and MEC respectively). The MEC receives its main inputs from brain regions implicated in spatial navigation, and conveys allocentric information that will be incorporated into multisensory assemblies in the EC. The MEC receives neocortical input via the postrhinal cortex, including a large input from retrosplenial cortex (Dubanet and Higley, 2024). It is in this area that grid cells are found, with receptive fields organized in hexagonal patterns, tiling the space where the mouse evolves (Fyhn et al., 2004). The LEC on the other hand, brings both egocentric and temporal information into the hippocampus (Tsao et al., 2018; Wang et al., 2018). The LEC receives neocortical input via the perirhinal cortex, in addition to direct input from olfactory bulb (Price, 1973). Beyond the division between ego- and allocentric function, the existence of a clean binary distinction between LEC and MEC is suggested by the segregation of the terminal zones of both regions in the molecular layer of the DG, with inputs from MEC and LEC occupying two distinct areas of the molecular layer (the middle one third for the terminal zone of MEC, and the outer one third for the terminal zone of LEC). Although MEC and LEC seem to be coherent entities with distinct properties at multiple levels, finer subdivisions of each region, clear at the cytoarchitectonic levels both in rodent and in primate brains, have been extensively described.

A few years ago, we undertook the first molecular profiling of principal neurons from layer IIa of EC (Roussarie et al., 2020). We used cell-type specific ribosomal profiling to obtain deep profiles of this neuronal population, at the expense of a single-cell view necessary to grasp the diversity of neuron types of layer IIa in various subregions of EC. Previously a bulk RNAseq study of MEC coupled with Allen Institute *in situ* hybridization (ISH) image analysis yielded clues into dorsoventral heterogeneity of the region, as well as lamina-specific gene expression patterns (Ramsden et al., 2015). Whole-brain single-cell atlasing efforts have captured to various extents the neurons from EC, but the description of EC subregions, and their molecular characteristics is still missing. Yet this is a necessary endeavor to understand the underpinnings of EC functional heterogeneity, for instance why neurofibrillary tangles first appear preferentially in specific EC areas (Braak and Braak, 1995; Llamas-Rodríguez et al., 2022). In this study, we profile mouse EC using single-nucleus RNAseq (snRNAseq) and highlight transcriptional types of neurons in the area. Focusing on layer IIa, we identify axes of heterogeneity within the ECIIa population. In parallel macaque EC studies, we also show that neuron subtypes in ECIIa seem to be largely conserved between rodents and primates. By defining genes enriched across distinct ECIIa subdivisions, this study provides a foundation for understanding their specialized functions and differential vulnerability.

## Methods

### Mice

Experiments with mice were approved by the Rockefeller University Institutional Animal Care and Use Committee (IACUC protocol #17039) or by the Boston University Institutional Animal Care and Use Committee (IACUC protocol #202100000007). All mice were maintained on a 12 h dark/light cycle and provided with food and water *ad libitum*. Mice used for single-nucleus RNAseq were 10-14 month old females carrying various apolipoprotein E (ApoE) alleles: mouse ApoE (1 mouse) – C57Bl/6J mouse from Jackson lab stock #00664; human ApoE E2 allele (3 mice) – Taconic stock #1547; human ApoE E3 allele (3 mice) – Taconic stock #1548; human ApoE E4 allele (4 mice) – Taconic stock #1549; mouse ApoE knockout (2 mice) – Taconic stock #APOE. At baseline, these mice with various alleles of ApoE are not expected to present differences in brain structure or function at baseline in the absence of amyloid or tau pathology. We also did not find any difference in the normative cellular taxonomy of the EC (see below and Supplementary Fig. 3. For ISH experiments 3-4 month-old male and female C57Bl/6J mice from Jackson lab were used (stock #000664).

### Single-nucleus RNAseq

To perform snRNAseq, we adapted a nucleus isolation protocol from (Jordi et al., 2013). Briefly, we slightly anesthetized mice with CO_2_ before decapitating them. We quickly dissected entorhinal cortices on a cold plate and placed the brain pieces in a chilled 1x Hank’s balanced salt solution containing 2.5 mM HEPES buffer at pH 7.4, 35 mM glucose, 4mM NaHCO_3_ and 5 mg/ml of actinomycin D to inhibit *de novo* transcription during the isolation, as in (Wu et al., 2017). We then placed the 2 entorhinal cortices of each mouse in 1 mL lysis buffer (20 mM HEPES buffer, 5mM MgCl_2_, 150 mM KCl, and 0.5mM DTT, 40 units/mL RNasin and 20 units/mL Superasin to inhibit RNases; we also added 5 mg/mL actinomycin D to this buffer) in a Kimble-Kontes 4 mL glass vial with a Teflon pestle, and lysed the samples with a motorized homogenization at 300 rotation per minute (rpm), followed by 12 strokes up and down the sample at 900 rpm. The lysate was then spun down at 500 x g for 10 minutes at 4C and the nucleus pellet resuspended in 29% iodixanol (D1556 from Sigma) solution containing 150 mM KCl, 5 mM MgCl_2_, 20 mM Tricine – KOH (pH 7.8), 5 mg/mL actinomycin D, 40 units/mL RNasin and 20 units/mL Superasin). Nuclei were then spun down for a total of 30 minutes at 10,000 x g at 4C, stopping the centrifugation every 10 minutes to remove the layer of myelin floating at the surface of the iodixanol using a pipet tip. After centrifugation, the supernatant was discarded and the nuclei resuspended in resuspension buffer (1% bovine serum albumin in phosphate buffered saline supplemented with 5 mg/mL actinomycin D, 40 units/mL RNasin and 20 units/mL Superasin and 10 mM DyeCycle Ruby (V10309) to stain nuclei). They were then sorted, selecting for single nuclei using DyeCycle Ruby. Nuclei were then sorted in resuspension buffer without DyeCycle Ruby. Nuclei were counted with trypan blue and a Countess cell counter, and loaded into a Chromium Controller. Sequencing libraries were synthesized using the 10x 3’ Gene Expression assays v2 and sequenced using a NovaSeq S2 sequencer with 150bp paired-end reads at the Rockefeller University Genomics Resource Center.

### Single-nucleus RNAseq Data analysis

#### Cell subtype identification

Raw fastq sequences were first aligned to the reference genome GRCm39-2024-A (downloaded from the 10x website) with cellranger-8.0.1 using default settings and intronic reads included. Count matrices for individual sample files were then imported into Seurat, which was used also for subsequent analyses (v5.4.0). Doublets were detected using Scrublet (Wolock et al., 2019) with an expected doublet rate of 0.8% and removed from the dataset. Individual samples were then merged into one dataset. The percentage of reads contributed by mitochondrial RNA was calculated for each nucleus and nuclei with more than 5% mitochondrial RNA reads were excluded. Additionally, nuclei with less than 400 genes (nFeature_RNA) detected were filtered out. We then log normalized gene counts, scaled data regressing number of UMIs and number of genes per nuclei (nCount_RNA and nFeature_RNA), selected the top 2,000 variable genes using the “vst” method, and ran a principal component analysis with 50 dimensions. For the general clustering, we clustered at a resolution of 0.7. To find markers enriched in the different clusters, we used the FindAllMarkers function with the default Wilcoxon test. We then subclustered inhibitory and excitatory neurons using 50 dimensions and resolutions of 0.3 and 0.7 respectively. After exclusion of non-EC excitatory neuron clusters (see results section), we performed a final clustering of excitatory neurons from EC with 50 dimensions and a resolution of 0.7. To investigate the heterogeneity of ECIIa neurons, we subclustered nuclei from LEC, CEnt and MEnt IIa neurons using 50 dimensions and a resolution of 0.1.

#### Cross-correlational analysis of subclusters

To evaluate correlations between subclusters, we averaged, for the most variable genes, scaled gene expression across all nuclei of the subcluster and then calculated for each pair of subclusters the Pearson correlation across these genes. We then corrected p-values with the Benjamini-Hochberg multiple testing correction. Hierarchical clustering was performed on both rows and columns of the Pearson correlation matrix using the ComplexHeatmap package in R. Dendrograms were generated utilizing Euclidean distance and the complete linkage clustering algorithm.

#### Identification of genes enriched in ECIIa subtypes

To identify genes differentially expressed between ECIIa neurons and all other excitatory neurons from EC, we merged clusters final_exc2, final_exc6 and final_exc10 from our final EC excitatory neuron clustering on the one hand, and all other clusters from the final_exc clustering on the other hand and performed pseudobulk differential gene expression analysis using DEseq2. For the gene set enrichment analysis, fast pre-ranked gene set enrichment analysis (FGSEA) (Korotkevich et al., 2021) was run from the full differential expression results. Genes were ranked from highest to lowest avg_log2FC from the entire list of results. No differential-expression significance cutoff was applied before creating the ranked list. We used the Mouse MSigDB M5 Gene Ontology collection, containing Biological Process, Cellular Component, and Molecular Function gene sets together, as well as the M2 WikiPathways collection. The GMT file was m5.go.v2026.1.Mm.symbols.gmt, using Mouse/MGI gene symbols from Mouse MSigDB v2026.1.Mm. Gene sets were loaded with fgsea v1.36.2 using fgsea::gmtPathways, and enrichment analysis was performed with fgsea::fgsea using the gene list ranked by avg_log2FC. Gene sets were restricted to sizes of 15–500 genes. A random seed was set before FGSEA. Pathways with Benjamini–Hochberg adjusted P value (FDR) < 0.25 were treated as significant for the filtered result tables and summary plots.

#### MapMyCells

To identify CCF supertype labels for all our excitatory neurons, we used the online interface of the MapMyCells tool, uploading the count matrix for cleaned up excitatory dataset, using the whole mouse brain reference CCN20230722 with hierarchical mapping. We then downloaded the MapMyCells output and matched supertype labels and final_exc cluster identity for all our cells. We calculated the percentage of identified supertype labels for each final_exc cluster.

#### Trajectory analysis

To test if subtypes of ECIIa neurons form a continuum or highly discrete populations we performed trajectory analysis on the clustered ECIIa dataset. The Seurat object was converted to a SingleCellExperiment (v. 1.32.0) while preserving the original cluster assignments, variable features and existing PCA embeddings (first 20 principal components). Trajectory inference was performed using Slingshot on the existing PCA (Street et al., 2018). Slingshot (v. 2.18.0) was run using the original clusters, without specifying a starting/root cluster, allowing the trajectory structure to be inferred from the cluster relationships in PCA space. Three lineages were identified: lineage 1 (ECIIa_7 → ECIIa_2 → ECIIa_4 → ECIIa_8 → ECIIa_1 → ECIIa_0), lineage 2 (ECIIa_7 → ECIIa_2 → ECIIa_4 → ECIIa_8 → ECIIa_1 → ECIIa_6) and lineage 3 (ECIIa_7 → ECIIa_2 → ECIIa_4 → ECIIa_8 → ECIIa_3 → ECIIa_5). For each nucleus, Slingshot pseudotime values and lineage-specific curve weights were retained for downstream analysis. Pseudotime-associated changes were then modeled using tradeSeq (v. 1.24.0) following the strategy developed by (Berge et al., 2020) . Briefly, the variable features already stored in the original Seurat object were used for modeling. Generalized additive models were fitted using tradeSeq::fitGAM with the default settings using slingshot pseudotime and lineage weights. The tradeSeq associationTest was then used to test whether gene expression changed significantly as a function of pseudotime. Multiple-testing correction was performed using Benjamini-Hochberg false discovery rate (BH FDR). Genes with FDR < 0.05 were considered significantly associated with pseudotime. Genes were also ranked by FDR for exploratory visualization of the most strongly dynamic genes. To plot expression of the genes as a function of pseudotime, we generated fitted expression profiles using tradeSeq predictSmooth with the default grid of 100 pseudotime points. These fitted profiles were used to determine the pseudotime position of maximum fitted expression (peak_index, calculated from the raw fitted predictSmooth expression values, rather than from z-scored values). Individual gene expression variation across pseudotime was represented with TradeSeq plotSmoothers plots showing the log fitted expression trends as a function of lineage-specific Slingshot pseudotime, with individual cells colored according to their original Seurat cluster. We also generated lineage-specific heatmaps with genes as rows, 100 fitted pseudotime positions as columns and tradeSeq fitted expression values as values. For visualization, fitted expression was z-scored independently for each gene across pseudotime. Therefore, heatmap colors represent expression relative to that gene’s own mean fitted expression rather than absolute expression differences between genes. Genes were ordered according to the pseudotime position of their maximum fitted expression, producing an early-to-late peak ordering. To relate the continuous fitted expression trajectories back to the original Seurat clusters, the heatmap columns were annotated with the dominant original cluster at each pseudotime position. For each lineage, cells with non-missing pseudotime were mapped onto the same fitted pseudotime grid used by predictSmooth. Where sufficient cells were available, cells with a lineage curve weight of at least 0.5 were preferentially used for this annotation. If fewer than 25 cells passed this threshold, all cells with valid pseudotime for that lineage were used instead. Cell pseudotime values were rescaled across the lineage and assigned to the corresponding fitted pseudotime bins. Within each bin, the most frequent original Seurat cluster was assigned as the cluster annotation. If a pseudotime bin contained no cells, its annotation was filled using the nearest pseudotime bin containing cells. This produced a continuous cluster annotation bar across the heatmap. The cluster annotation is therefore a descriptive mapping of the continuous Slingshot trajectory back onto the original clustering and does not represent a new clustering analysis.

### In Situ Hybridization (ISH)

*Mouse.* For mouse ISH and ISH-IF co-detection, we used the RNAscope fluorescent v2 kit (Advanced Cell diagnostics) on fixed frozen tissue, following, for most steps, the manufacturer’s manual. Briefly mice were anesthetized with a mixture of ketamine (90mg/kg) and xylazine (10mg/kg) and, once deeply anesthetized, transcardially perfused with 1x PBS, followed by 4% PFA in 1x PBS. The brain was then dissected and postfixed for 24 hours at 4°C in 4% PFA in 1x PBS. The brain was then cryoprotected in 10%, 20% and 30% sucrose in 1x PBS, before being embedded in OCT and frozen on dry ice. Before sectioning, the brain was equilibrated to −20°C, sectioned at a thickness of 10 µm, and mounted on superfrost slides that were left to dry for 1-2 hours inside the cryostat before being stored at −80C. *ISH.* Selected slides were removed from the −80°C and washed in 1x PBS for 5 minutes. Slides were then baked for 30 minutes at 60°C (HybEZ Hybridization Oven) and washed in 1xPBS for 5 minutes before post-fixing the slides in pre-chilled 4% PFA in 1xPBS at 4°C 15 minutes. Following dehydration steps of 50%, 70%, 100% ethanol for 5 minutes at room temperature each dilution, slides were air dried. They were then incubated for 10 minutes at room temperature with RNAscope Hydrogen Peroxide. After 2 washes in water, slides were submerged into 94-98°C RNAscope 1x Target Retrieval solution for 5 minutes. Immediately after target retrieval, slides were washed in water, transferred to 100% ethanol for 3 minutes and air dried. Once the slides were dry, a hydrophobic barrier was drawn around the tissue (ImmEdge Pen, Vector laboratories) and slides were dried overnight at room temperature (RT). RNAscope Protease III was added on each piece of tissue before incubating for 30 minutes at 40°C. In the meantime, probes were warmed for 10 minutes at 40°C before preparing the probe mix (1:50 of C2-C4 probes diluted in C1 probe). After 2 washes in water, slides were incubated with probes for 2 hours. After probe hybridization, the RNAscope protocol was followed, using opal dyes to label the different probes (1:1,500 dilution of each dye except 1:750 for Opal 780). The slides mounted with Prolong Gold antifade reagent. Slides were allowed to dry overnight before imaging using the Vectra Polaris slide scanner. *ISH-RELN IF co-detection.* For the co-detection of mRNA and protein, selected slides were removed from the −80°C and washed in 1x PBS for 5 minutes. Slides were then baked for 30 minutes at 60°C (HybEZ Hybridization Oven) and washed in 1xPBS for 5 minutes before post-fixing the slides in 4% PFA in 1xPBS for 1 hour at RT. Following dehydration steps of 50%, 70%, 100% ethanol for 5 minutes at room temperature for each dilution, slides were air dried. They were then incubated for 10 minutes at room temperature with RNAscope Hydrogen Peroxide. After 2 washes in water, slides were submerged into 94-98°C RNAscope 1x co-detection Target Retrieval solution for 5 minutes. Immediately after target retrieval, slides were washed twice in water, and once in PBS-T (1x PBS with 0.1% Tween-20, pH 7.2-7.4). A hydrophobic barrier was drawn around the tissue, and the RELN (PA5-47537) primary antibody was incubated with the tissue overnight at 4°C at a 1:50 dilution in RNAscope co-detection antibody diluent. The following day, slides were washed three times in PBS-T and fixed again in 10% neutral buffered formalin for 30 minutes at RT. After post-primary fixation, slides were washed four times in PBS-T and digested with RNAscope Protease III for 30 minutes at 40°C. In the meantime, probes were warmed for 10 minutes at 40°C before preparing the probe mix (1:50 of C2-C4 probes diluted in C1 probe). After 2 washes in water, slides were incubated with probes for 2 hours. After probe hybridization, the RNAscope protocol was followed, using opal dyes to label the different probes (1:1,500 dilution). For detection with Opal 780, only the incubation with HRP-digoxigenin was performed at this point. After the last wash in RNAscope wash buffer, the secondary antibody to detect RELN (Alexa-488 coupled donkey anti-goat polyclonal antibody) was incubated with the sections for 30 minutes at RT (diluted 1:300 in RNAscope co-detection antibody diluent) and slides were washed twice in PBS-T. Lastly, slides were incubation with Opal 780 for 30 minutes at RT before 2 washes in PBS-T, incubation with RNAscope DAPI, and mounting with prolong gold reagent.

#### Macaque

Origin of tissue: 30µm hemisected coronal sections were obtained from a 9-year old male subject from the archival collection of 4% PFA perfused Macaca mulatta brain tissue from Dr. Rosene at Boston University Chobanian and Avedisian School of Medicine. Sections are stored in cryoprotectant buffer (0.1M phosphate buffer, with 15% glycerol and 1% PFA) at −80°C. A piece of tissue containing the EC was selected from fixed free floating 30µm Macaca mulatta hemisected sections. After a wash with 1x PBS, free floating tissue was placed in a 24 well plate with RNAscope Hydrogen Peroxide for 10 minutes at room temperature with a gentle rock. The tissue was then washed with water twice before mounting it on Superfrost plus slides in 1x PBS. After drying of the section for 5 minutes at room temperature, each slide was dipped in 100% ethanol a few times and allowed to dry briefly. Once dry, slides were fully submerged into 92-95°C hot RNAscope 1x Target Retrieval for 5 minutes, washed in water, dipped in 100% ethanol and dried briefly before drawing a hydrophobic barrier around the tissue. The sections were then digested with RNAscope Protease Plus for 30 minutes at 40°C. In the meantime, primary probes were warmed for 10 minutes at 40°C before preparing a probe solution (1:50 of C2-C4 probes diluted in C1 probe). After 2 washes in water, slides were incubated with primary probes for 2 hours. Slides were washed twice in RNAscope Wash Buffer in a wash container followed by a 30 minute incubation with RNAscope multiplex FL v2 Amp1, a 30 minute incubation with RNAscope multiplex FL v2 Amp2 and a 15 minute incubation with RNAscope multiplex FL v2 Amp3, washing twice in wash buffer between amplification steps. After amplification, slides were incubated with RNAscope multiplex FL v2 HRP-C1 to HRP-C4 in a repeated process consisting in opening the channel, adding the desired dye and closing the channel as follows: 15 minute incubation with HRP-C1, wash twice in wash buffer, 30 minute incubation with Opal reagent diluted 1:750 in RNAscope Multiplex TSA buffer, wash twice in wash buffer, and 15 minute incubation with RNAscope multiplex FL v2 HRP blocker. This process was repeated for each channel used and its designated Opal fluorophore color. After all channels have been incubated with its designated fluorophore and the last 2 washes were performed, a few drops of RNAscope multiplex FL v2 DAPI were added on each piece (one at a time) for 30 seconds at room temperature. The slides were flicked to remove excess liquid and a coverslip was added with Prolong Gold antifade reagent. Slides were allowed to dry overnight before imaging using the Vectra Polaris slide scanner.

#### Probes

we used the following probes for ISH (from Advanced Cell Diagnostics): Mm-Rfx3 (313371), Mm-Erbb4 (318721), Mm-Tenm3-C1 (411951), Mm-Kirrel3-C1 (463651), Mm-Lef1-C2 (441861-C2), Mm-Gpc5-C2 (442831-C2), Mm-Cdh8-C2 (485461-C2), Mm-Reln-C3 (405981-C3), Mm-Il1rapl2-C3 (552341-C3), Mm-Cdh18-C3 (1183161-C3), Mm-Grik1-C3 (438771-C3), Mm-Dcn-C4 (413281-C4), Mm-Adam33-C4 (552321-C4), Mm-Erbb4-C4 (318721-C4), Mm-Reln-C4 (405981-C4), Mm-Tenm3-C4 (411951), Mmu-VWC2L-C3 (1246551), Mmu-GPC5-C2 (1095881-C2), Mmu-RELN-C4 (1181301-C4), Hs-TENM3-C1 (411941), Hs-DCN-C1 (589521), Hs-RELN-C2 (413051-C2), Hs-GRIK1-C3 (836311-C3), Hs-ERBB4-C4 (407831-C4).

## Results

To investigate molecular heterogeneity of EC we performed snRNA-seq using 10x genomics with nuclei isolated from dissected EC of 10-14 month-old female mice. After filtering high-quality single nuclei, we obtained 47,573 nuclei, clustered them, and identified all major cell types in the expected proportions using known marker genes, i.e. excitatory neurons (65.0 %, expressing *Slc17a6* or *Slc17a7*), inhibitory neurons (15.4 %, expressing *Gad1* and/or *Gad2*), astrocytes (7.5 %, expressing *Gja1*), oligodendrocyte progenitor cells (5.1 %, expressing *Pdgfra*), microglia (2.4 %, expressing *Siglech*), oligodendrocytes (2.0 %, expressing *Mag*), pericytes (0.9 %, expressing *Slc6a13* (Buizza et al., 2025)), fibroblast-like vascular cells (0.6 %, *Fxyd5* (Saunders et al., 2018)), endothelial cells (0.3 %, *Flt1* (Daneman et al., 2010)) and border-associated macrophages (0.2 %, expressing *F13a1* (Utz et al., 2020)) (Supplementary Fig. 1; Supplementary Table 1). We re-clustered all excitatory and inhibitory neurons.

### Inhibitory neuron type distribution within EC

The subclustering of EC inhibitory neurons and marker enrichment analysis for each cluster allowed us to identify all major types of inhibitory neurons previously described (Supplementary Fig. 2, Supplementary Table 2). We found interneurons originating from the medial ganglionic eminence: parvalbumin (*Pvalb*) interneurons (cluster inh_0), chandelier cells – *Pvalb*, *Pthlh* and *Vipr2* positive (inh_07), *Npy*-positive and *Npy*-negative somatostatin (*Sst*) interneurons (inh_01 and inh_05 respectively) and ivy cells - *Lamp5*/*Lhx6* co-expressing interneurons (inh_08) (Krienen et al., 2020). The *Npy*/*Sst* interneurons are likely to be Martinotti cells (Urban-Ciecko and Barth, 2016). We also found neurons from the caudal ganglionic eminence: *Vip*-expressing interneurons (inh_02), *Lamp5*/*Ndnf*-co-expressing interneurons (inh_03), *Sncg*-expressing interneurons (inh_04), *Ntng1*/*Ndnf*-co-expressing interneurons (inh_06), which correspond respectively to bipolar, neurogliaform, multipolar and retrohippocampal-cortical amygdala (RHP-COA) interneurons - as defined by the Allen Institute (Velthoven et al., 2025). While we predominantly profiled the EC, a minor proportion of starting material may have originated from neighboring neocortex, piriform and subiculum. A minority of these interneurons might thus originate from these areas.

### Excitatory neuron clustering

Excitatory neuron types are much more spatially heterogeneous than inhibitory neurons, with layer- and subregion-specific gene expression patterns. It is therefore possible to distinguish EC neurons from neurons in neighboring regions unavoidably co-dissected with EC. We first filtered our EC dataset by excluding neurons originating from regions other than EC: mining the Allen Institute mouse brain ISH dataset (Lein et al., 2007) for markers enriched in the initial clusters, we identified excitatory neuron types corresponding to all clusters, and proceeded to exclude the following: neurons from the cortical nucleus of amygdala (clusters general_3 and general_8), recognized by their expression of *Trh* and co-expression of *Trhr and Lypd1* respectively; posteromedial amygdalo-hippocampal transition area (cluster general_12 co-expressing *Htr4* and *Pde11a*); amygdalo-piriform transition area (clusters general_13 co-expressing *Htr2c* and *Tmem150c*); ventral subiculum (clusters general_22 and general_23 expressing *Sntb1* and *Bace2* respectively); endopiriform nucleus (cluster general_25 expressing *Spag16*), presubiculum (cluster general_27 expressing *Npsr1*) and dentate gyrus granule cells (cluster general_26, expressing *Prox1*) (Supplementary Table 1). After reclustering all other excitatory neurons and marker enrichment analysis (Supplementary Fig. 3 and Supplementary Table 3), we further excluded neurons from Piriform cortex (cluster exc_0 enriched in *Slco2a1* and cluster exc_14 enriched in *Erbb4* but without *Reln*), neurons from temporal association cortex layer II/III (cluster exc_7 enriched in *Naa11*), layer V (cluster exc_20 enriched in *L3mbtl4*), layer V/VI (cluster exc_19 enriched in *Oprk1*), and layer VI (cluster exc_17 enriched in *Hs3st5*).

### Entorhinal cortex excitatory neurons

We then performed a final round of reclustering of the cleaned-up dataset to obtain a full molecular picture of excitatory neuron types in EC (15,654 nuclei, Fig. 1). We verified that the various ApoE genotypes of the mice that we profiled did not seem to affect the normative cellular taxonomy of excitatory neurons of the EC (as demonstrated by the intermingling of genotypes for all transcriptomic clusters, see Supplementary Fig. 3). We obtained marker genes enriched in each of the subclusters (Table 1). Use of the mouse Allen Brain Atlas ISH data allowed us to identify the anatomical location of each subcluster. The cytoarchitectonic structure of EC was reflected, at the molecular level, by the existence of layer-specific molecularly defined types of neurons. In addition, the differences long known to exist between LEC and MEC were also clear at the transcriptomic level with distinct MEC and LEC clusters for several but not all excitatory neuron types: at the resolution chosen in Figure 1, external layers, the ones superficial to *lamina dissecans* were all LEC- or MEC-specific. With the Witter nomenclature, layer II is subdivided in layers IIa and IIb based on their expression of Reelin (*Reln*) and Calbindin (*Calb1*) respectively.

**Figure 1.**
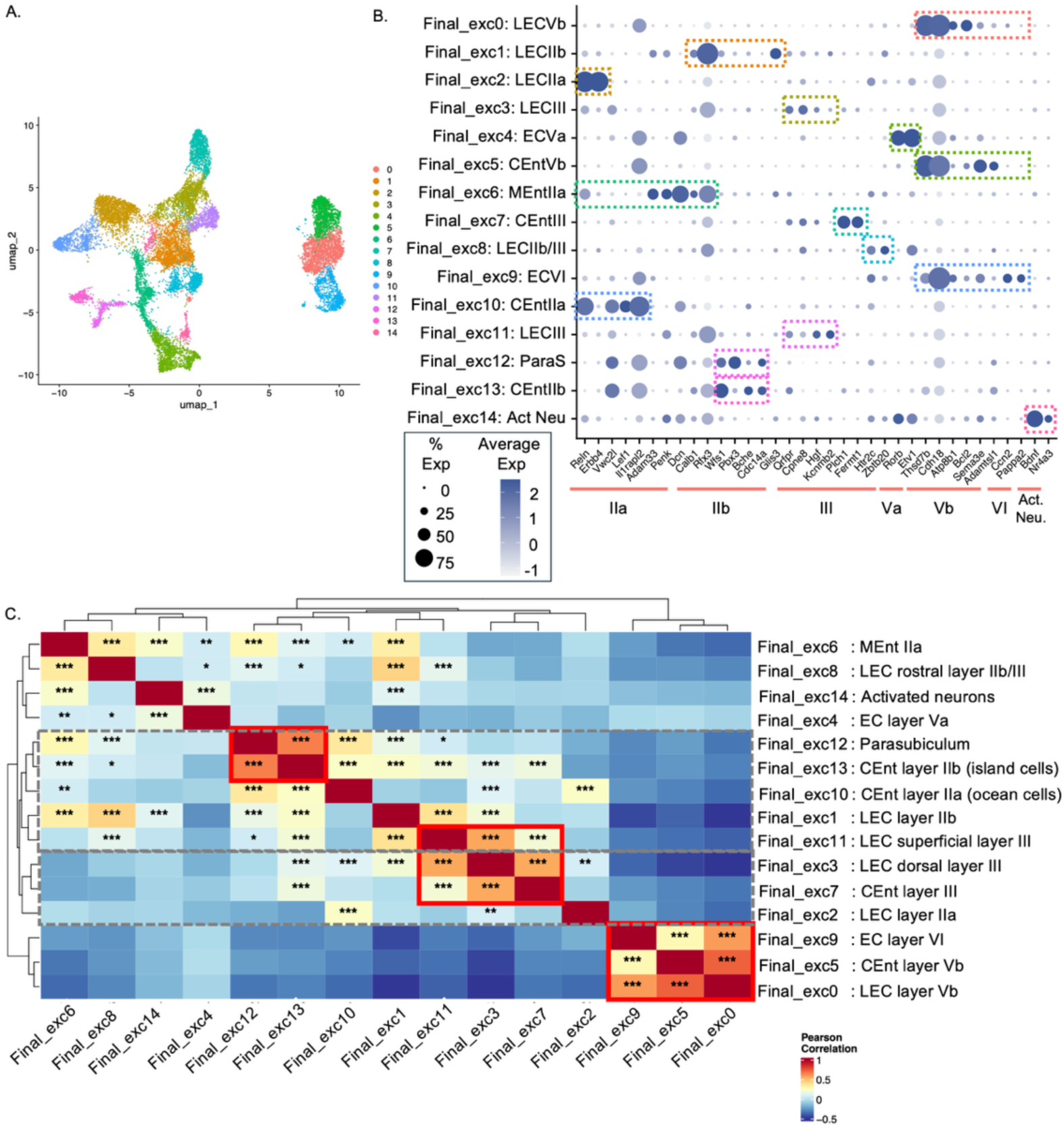
Clustering of all EC excitatory neurons. After identification of excitatory neurons, reclustering, exclusion of non EC excitatory neuron clusters, we performed a final clustering (final_exc). A. Uniform manifold approximation and projection (UMAP) visualization of the 15 different EC excitatory neurons clusters. B. Dot plots for excitatory neuron subtype-specific markers in each of the 15 final_exc clusters. Dots are colored by level of expression of the marker in the cluster, and their size correlates with the percentage of expression of the marker in the cluster. C. Pearson correlation between pairs of final_exc clusters. We also performed hierarchical clustering on all clusters. We outlined in red the groups of clusters showing highest correlations with one another: deep layers (Vb and VI – final_exc0, final_exc5, final_exc9); all layers III (CEnt, dorsal LEC, superficial LEC – final_exc3, final_exc7, final_exc11); CEnt layer IIb (island cells) and parasubiculum (final_exc12 and final_exc13). We outlined in gray the two branches of the hierarchical clustering corresponding to superficial layers of LEC and CEnt to show the complex relationship between these populations. LECIIa and the other *Reln*-expressing excitatory type LEC dorsaly layer III co-segregated with CEntIII, while CEntIIa, CEntIIb and LECIIb were found in a distinct branch. The cross-correlogram is colored by Pearson correlation coefficient. The significant correlations after Benjamini-Hochberg multiple testing correction are marked with asterisks.

We first identified *Reln*-expressing layer IIa neurons. Surprisingly, not two but three distinct clusters of excitatory neurons expressed *Reln* in EC – clusters final_exc2, final_exc6 and final_exc10 (overall these three clusters amounted to 3,385 nuclei, i.e. 21.6% of EC excitatory neurons) (Table 1 and Fig. 1). These three populations correspond respectively with layer IIa neurons from LEC (*Reln* and *Erbb4* co-expression, cluster final_exc2), from the caudomedial subdivision of entorhinal (CEnt) (*Reln* and *Vwc2l*, *Lef1* or *Il1rapl2* co-expression, cluster final_exc10), and from the medioventral subdivision of entorhinal (MEnt) (*Reln* and *Adam33*, *Penk* or *Dcn* expression, cluster final_exc6), which we found to be different enough from CEnt layer IIa to form their own cluster. This is in agreement with the whole mouse brain taxonomy from Allen Brain Cell (ABC) atlas (reference CCN20230722), which also distinguishes ENTl (lateral, our LEC), ENT-po (posterior, our CEnt) and ENTmv (medioventral, our MEnt). CEnt and MEnt, which together are thought to constitute the MEC, have previously been recognized as different in cytoarchitectonics studies (Groen, 2001; Groen et al., 2003), and are distinct in the Paxinos atlas (Paxinos and Franklin, 2012). But work from electrophysiologists and behavioralists traditionally groups them together under the overarching MEC category. Transcriptomic data from our group and the Allen Institute suggest that the superficial *Reln*-positive cell layer is best subdivided into three rather than two cell populations. To clearly visualize these three subdivisions in mouse brain sections, we performed a multiplexed ISH for *Erbb4*, *Lef1*, *Adam33* and *Reln* on horizontal brain sections, and classified neurons based on their expression of *Reln* and *Erbb4*, *Reln* and *Lef1* or *Reln* and *Adam33* (Fig. 2A). Layer IIa of CEnt is the only population visible on the dorsal-most sections. On intermediate sections, the cells in the medial-most half of the layer, where it is most curved, are part of CEnt, while the cells in the lateral/rostral half of the layer are in the LEC. In more ventral sections, LECIIa becomes more prominent, and CEntIIa gives way to MEntIIa on the ventral most sections. We also successfully detected CEntIIa cells with a probe directed against *Il1rapl2*. The border between LECIIa and CEntIIa is distinguishable solely with a staining for Reelin (either *Reln* mRNA as in Fig. 2A or RELN protein as in Fig. 2C) since Reelin is expressed at higher levels in the LEC, and the layer is thinner and denser than in CEnt. Supplementary Fig. 4 shows the position of LECIIa, MEntIIa and CEntIIa from the Allen mouse brain common coordinate framework (CCF) ABC atlas (with supertypes 0042 corresponding to LECIIa, 0066, 0067, 0068 to MEntIIa and 0051, 0052, 0053 and 0054 to CEntIIa).

**Figure 2.**
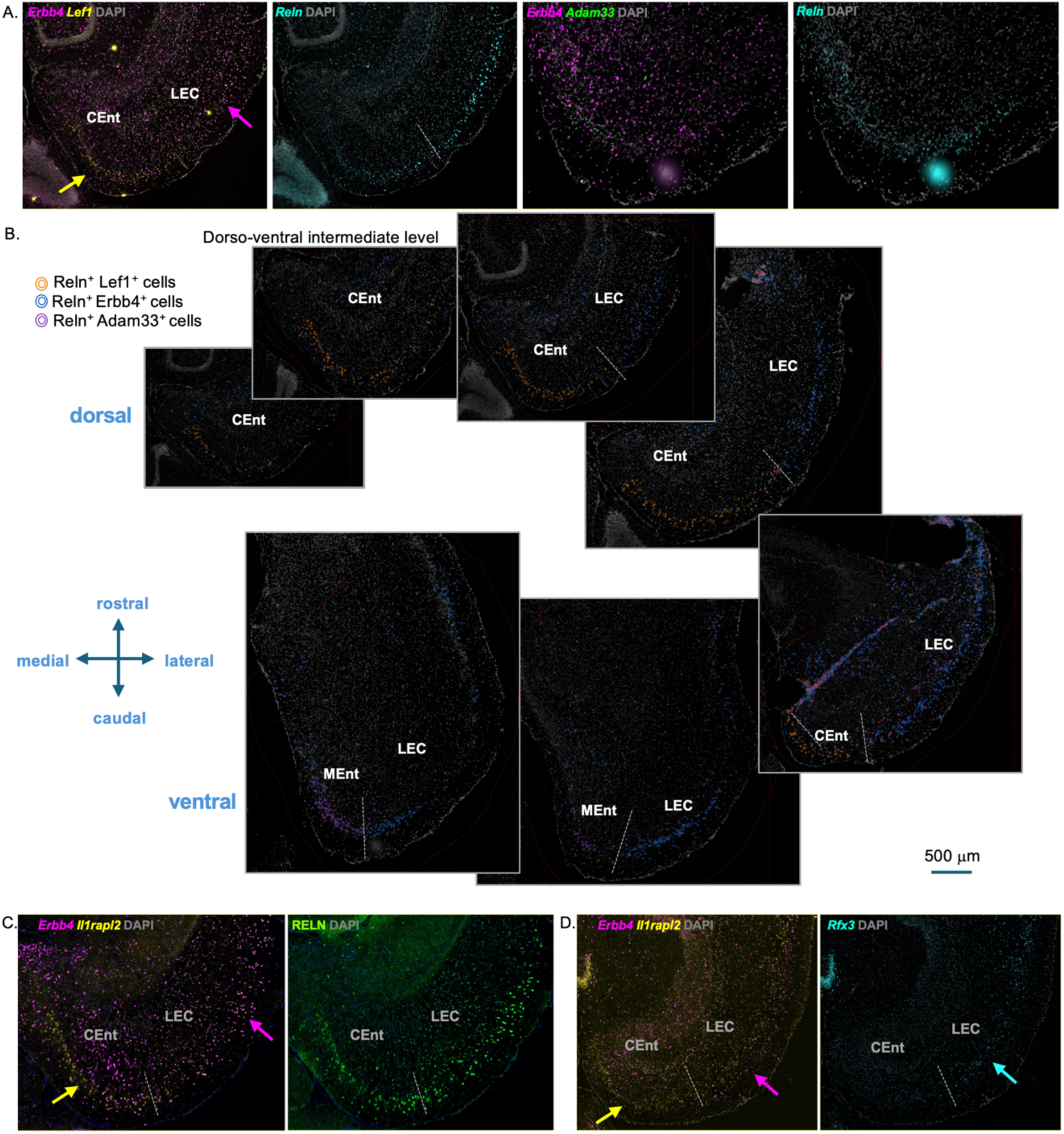
Identification of CEntIIa, LECIIa and MEntIIa in horizontal sections of a 3-month-old C57Bl/6J female mouse brain. A. ISH detection of *Erbb4*, *Lef1*, *Reln* and *Adam33*. Nuclei were counterstained with DAPI. B. Classifiers detecting cells expressing *Reln* and *Lef1* (CEntIIa, orange), *Reln* and *Erbb4* (LECIIa, blue) and *Reln* and *Adam33* (MEntIIa, purple). Cells were detected using DAPI, classified based on their expression of Reln, and then further classified based on their expression of the other three markers. Horizontal sections arranged from dorsal-most to ventral-most and spanning the entire extent of the EC are shown. C. co-detection of *Il1rapl2* and *Erbb4* by ISH and RELN by IF showing enrichment of CEntIIa neurons in Il1rapl2. D. ISH detection of *Il1rapl2*, *Erbb4* and *Rfx3*. The magenta arrows point to ECIIa cells expressing *Erbb4*. Yellow arrows point to CEntIIa cells expressing Lef1 (A) and Il1rapl2 (C and D). The cyan arrow points to LECIIb cells expressing *Rfx3*. Of note, all interneurons express *Erbb4*, so scattered cells can be found throughout EC expressing it. Similar results were obtained in coronal sections of another 3-month old C57Bl/6J female mouse.

To find the cluster correlates of layer IIb neurons we searched for clusters with significant enrichment of Calbindin (*Calb1*). Strongest enrichment was found in final_exc1 (Table 1). This cluster is also where *Rfx3* is most significantly enriched. ISH for *Rfx3* demonstrated that final_exc1 were neurons located immediately beneath *Erbb4*-stained layer IIa in LEC specifically (Fig. 2D), i.e. layer IIb as described by Witter(Witter et al., 2017) (this ISH additionally highlights the border between LEC and CEnt – with conspicuously lower *Rfx3* expression in CEnt). On the other hand, both Witter and Tonegawa have described that IIa and IIb excitatory neurons were not found in distinct laminae in MEC like they are in LEC. At least in the CEnt subdivision, Kitamura *et al* found that patches of Reln cells (they referred to them as “ocean cells”) alternate with patches of *Wfs1*-positive cells (referred as “island cells”) (Kitamura et al., 2014). We found significant *Wfs1* enrichment in clusters final_exc1 (LECIIb), final_exc12 and final_exc13 (Fig.1A). To determine which of these two latter clusters corresponded to “island cells” or CEnt layer IIb neurons, we searched for genes differentially expressed between them. *Pbx3* was only expressed in final_exc12. From Allen ISH data, the parasubiculum is the main group of neurons in the area with high Pbx3 expression (Supplementary Fig. 5A). On the other hand, *Bche* was expressed in island cells but not in parasubiculum (Supplementary Fig. 5A). We conclude that final_exc12 represents parasubicular neurons, final_exc13 island cells or CEntIIb, and that these two cell types present striking molecular resemblance with one another, as suggested by previous literature (Sammons et al., 2019). Calb1 is enriched in CEntIIb (not in parasubiculum final_exc12, consistent with previous findings (Sammons et al., 2019)), but also in final_exc3 and final_exc11, and is therefore a less specific IIb marker than *Wfs1*. Along with *Wfs1*, final_exc12 is characterized by high enrichment of *Cdc14a*, which is specific to islands/CEntIIb and not expressed in LECIIb. *Glis3*, on the other hand, is a specific marker for LECIIb – cluster final_exc1, not expressed in islands/CEntIIb –final_exc13. Lastly, the MEnt subdivision, at least at this resolution did not seem to include distinct subtypes of cells for IIa and IIb. Staining for markers of IIa and IIb neurons (Reln and *Rfx3*) reveals two layers in the LEC that essentially merge into one thick layer of *Reln*-*Rfx3* co-expressing neurons at the level of the MEnt (Supplementary Fig. 5B). The snRNAseq data suggests that they are also significantly enriched in *Calb1*. Taken together, we find that IIa and IIb neurons from known EC subdivisions have distinct molecular correlates, and we single out markers to identify them, beyond the already known *Reln* (IIa) and *Wfs1* (IIb): *Erbb4* for LECIIa, *Il1rapl2* and *Lef1* for CEntIIa, *Glis3* and *Rfx3* for LECIIb, *Cdc14a* and *Bche* for CEntIIb; both *Reln* and *Rfx3* for MEnt where IIa and IIb seem to form a single layer. Temporoammonic CA1 projecting neurons were previously shown to be the *Wfs1*-expressing neurons in the MEC and LEC (Kitamura et al., 2014; Ohara et al., 2019; Delpech et al., 2021). They are therefore likely to be LECIIb and CEntIIb (final_exc_1 and final_exc_13).

Further matching snRNAseq markers with Allen ISH expression profiles, we found 3 clusters of neurons corresponding to the layer immediately beneath layers IIa and IIb, i.e. layer III: in LEC, final_exc3 (dorsal, including around the rhinal fissure, more abundant in caudal areas, with enrichment of *Qrfpr* and *Cpne8*) and final_exc11 (superficial to final_exc3 and more abundant in rostral and ventral areas, with enrichment of *Hgf* and *Kcnmb2*); in MEC, final_exc7 (enriched in *Plch1* and *Fermt1*). A last cluster corresponding to external layers, final_exc8, enriched in *Htr2c* and *Zbtb20* seems to be found in the most rostral parts of LECIIb/III. Final_exc3 is the only excitatory neuron population with *Reln* expression, outside of layer IIa. This is likely the homologue of the human deep layer III which also shows RELN expression (Rodriguez-Rodriguez et al., 2025).

Of note, *Cux2*, a marker for neocortex layer II, is broadly enriched in all EC excitatory neuron types from layer IIb and III (final_exc1, final_exc3, final_exc7, final_exc11, final_exc13), except in the most rostral LECIIb/III population.

Deep to the *lamina dissecans*, we found a transcriptional landscape with less LEC/MEC heterogeneity. While layer Vb neurons (Thsd7b and Cdh18 positive cells) were still divided into a LEC (*Atp8b1* and *Bcl2* positive, final_exc0) and a MEC (*Sema3e* and *Adamtsl1*, final_exc5) subset, neurons from LEC and MEC in layers Va (*Rorb* and *Etv1* positive, final_exc4) and VI (*Ccn2* and *Pappa2* positive, final_exc9) did not form separate clusters. The layer Vb clusters, significantly enriched in *Bcl11b* and *Pcp4* (Table 1) correspond to the cells receiving CA1 inputs (Ohara et al., 2018).

Cluster final_exc14, a cluster enriched in layer Va markers like *Rorb* and *Etv1* but distinct from the main layer Va cluster final_exc4, was not obviously identifiable as an anatomical subtype within the layer. On the other hand, we noticed that many final_exc14 markers were traditionally linked with neuronal activity. Indeed, we found that are a gene set of delayed primary response genes, genes expressed as a result of sustained neuronal activity, were significantly enriched in the cluster (gene set from (Tyssowski et al., 2018), Wilcoxon rank-sum test p-value < 2.2 x 10^-16^). Interestingly, this cluster had similarity both with layers IIb, III and Va neurons, suggesting that it might include three distinct neuron types with a shared activity-regulated program. To test this hypothesis, we subclustered final_exc14 (Supplementary Fig. 6 and Supplementary Table 4). We obtained subclusters with markers of layer Va (*Rorb*, final_exc14_subcluster0), LECIIb (*Rfx3*, final_exc14_subcluster1), MEntII (*Car12* and *Penk*, final_exc14_subcluster 2) and LECIII neurons (*Fign*, final_exc14_subcluster 3). Notably, *Lrrk2*, the genetic susceptibility factor for Parkinson’s disease (Rocha et al., 2022) was enriched in final_exc14_subcluster3, but not in the main LECIII cluster, suggesting that it might be a LECIII-specific activity-induced gene. Overall, final_exc14 corresponds to various neuronal populations from EC: its molecular profile results from the superimposition of both a neuron-type specific program and a neuronal activity-induced program similar across these neuron types. We did not find an equivalent activation signature in other excitatory neuron populations, although there are individual neurons, for instance in LECIIa, with expression of delayed primary response genes like *Nr4a3* or *Bdnf*. Either these cells are too rare to form a distinct cluster, their activity-induced signature is less prominent than their neuron-type specific transcriptional program, or there are various expression programs in active IIa cells that prevent them from co-clustering.

### Visualization of the global laminar structure of EC

To visualize the different cell layers simultaneously, we performed multiplexed co-detection IF/ISH for a set of markers that covers some of the main EC layers: RELN protein (layer IIa), *Rfx3* (layer IIb), *Cdh8* (layer III), *Cdh18* (layer Va) and *Dcn* (layer Vb/VI). Supplementary Fig. 7 shows the raw ISH signal (top series of images), as well as the result of classifier detecting expression of the various markers (bottom series of images).

As an alternative approach to grasp the molecular anatomy of the region, we sought to leverage the Allen Institute spatial transcriptomic atlas and the CCF. We first used the Allen MapMyCells tool (Yao et al., 2023), to determine the CCF supertype of each of our snRNAseq nucleus (using the whole mouse brain reference CCN20230722), as well as the percentage of each supertype representation in each of our final_exc clusters (Supplementary Table 5). Leveraging the ABC atlas, we then plotted our final_exc clusters onto coronal sections of the atlas, grouping for each final_exc cluster all supertypes representing more than 15% of cells of the cluster. For supertypes present in several clusters, we attributed them the final_exc label where they have highest representation. Supplementary Fig. 8 shows the rostrocaudal series of coronal sections.

### Correlation between EC excitatory neuron subtypes

The taxonomic separation of all cells into neuron types obviously depends on the granularity with which the cells are compared, i.e. the resolution of our clustering. To determine more systematically the molecular similarities between the above-mentioned excitatory neuron subtypes, we calculated pairwise Pearson correlations for whole-genome gene expression levels with each pair of clusters.

We then performed hierarchical clustering of the clusters. The results are presented in Fig. 1C. Strikingly, we find correlation between excitatory subtypes in the most internal layers (II-Va), between excitatory subtypes in the most external layers (Vb and VI – red frame in Fig. 1C), but not between these two groups. In particular, there is very high correlation between Vb in LEC and in MEC. That the only deep population that diverges between LEC and MEC is actually relatively similar in both areas neurons confirms the molecular continuity of internal layers across the entire EC. On the other hand, both subregion and layer determine gene expression signatures of excitatory neurons from external layers. CEntIIa correlates well with LECIIa and MEntII. Dorsal LECIII (final_exc3), which is found in the same area as both LECIIa (in rostral parts of EC) and CEntIIa (in caudal parts of EC) correlates with both, in addition to correlating very highly with ventral LECIII and CEntIII (red frame in Fig. 1C). Layers IIb and III strongly correlated with one another, across LEC and CEnt. There was also very high correlation between parasubiculum and CEntIIb island cells, confirming the similarity that we mentioned earlier (red frame in Fig. 1C).

Altogether, this snRNAseq experiment paints the picture of a heterogeneous region, where the molecular makeup of excitatory neurons is determined by complex laminar architecture and regional subdivisions, especially above the *lamina dissecans*. Given the intricate connectivity between pyramidal cells from layer II-III and ECIIa neurons (reviewed in (Tukker et al., 2022)), one can hypothesize that these layers are constituted by a mosaic of unique local circuits.

### Molecular characteristics of ECIIa neurons

Layer IIa neurons are uniquely vulnerable to NFT formation, compared to excitatory neurons of other layers. With our snRNA-seq dataset we can start to search for the molecular features that could underlie their vulnerability. To that end, we pooled nuclei from all three ECIIa clusters (LEC, CEnt, MEnt) and compared their molecular profiles to those of nuclei from all other excitatory neuron types. We found 467 genes enriched in ECIIa vs. other neurons, and 640 genes impoverished there (at an adjusted p-value (padj) < 0.05) (Fig. 3A and Supplementary Table 6). We then performed gene set enrichment analysis (GSEA) on the differential expression results, using gene ontology (GO) pathways (Fig. 3B). We find significant enrichment of genes involved in cholesterol biosynthesis (padj = 6.0 x 10^-3^, normalized enriched score (NES) = 1.96). Many enzymes involved in cholesterol synthesis showed higher expression in ECIIa neurons compared to other neurons of the region, although the enrichment of each enzyme was modest (significant enrichment of Sqle, nominally significant enrichment of Dhcr7, Msmo1, Cyp51, Fdps, Hmgcr, Hmgcs1, Sc5d, Idi1). These enzymes span almost the entire mevalonate/cholesterol synthesis pathway from Acetyl-coA to cholesterol and suggests increased activity of the master regulator of cholesterol synthesis, SREBP2 (Horton et al., 2002), in ECIIa neurons. We indeed found modest enrichment of *Srebf2*, which encodes SREBP2, in ECIIa neurons (p-value = 0.03, log2FC = 0.16). Although the protein level of these enzymes is known to be heavily regulated at the posttranslational level (Sharpe et al., 2020), higher transcription rate in ECIIa neurons might indicate greater need for cell-autonomous synthesis of cholesterol compared to other excitatory neurons of the region.

**Figure 3.**
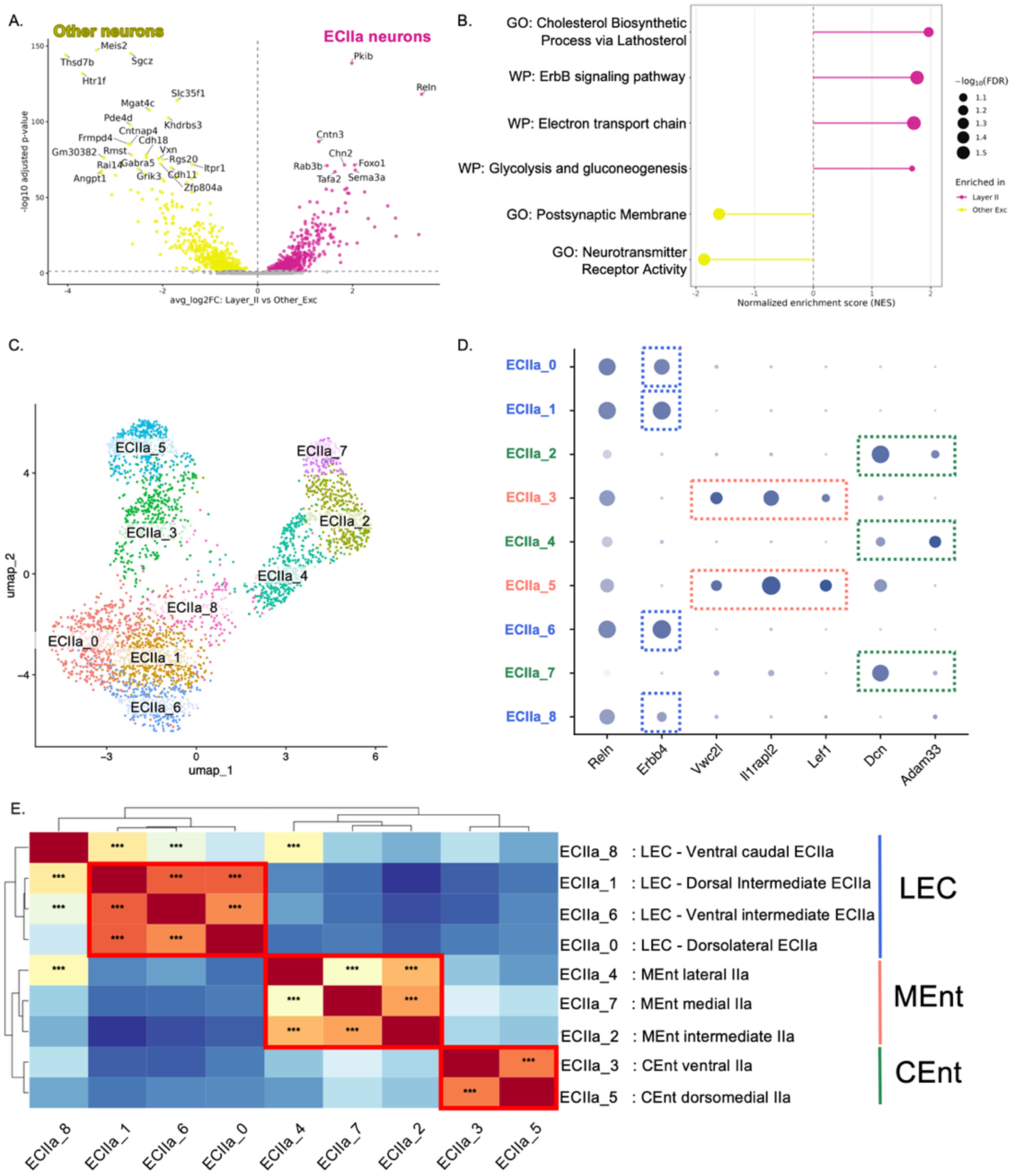
Characterization of ECIIa neurons. We analyzed genes differentially expressed between ECIIa neurons and other EC excitatory neurons (A-B). A. Volcano plot of differentially expressed genes. Gene expression pseudobulked across all ECIIa neurons (final_exc2, final _exc_6 and final_exc_10) was compared to pseudobulked gene expression across all other excitatory neurons using DESeq2. B. GSEA analysis of the differentially expressed genes using GO pathways and wikipathways (WP). We then subclustered all ECIIa excitatory neurons (C-E). C. UMAP visualization of the 9 ECIIa excitatory neurons clusters. D. Dot plots for ECIIa neuron subtype-specific markers in each of the 9 ECIIa clusters. Dots are colored by level of expression of the marker in the cluster, and their size correlates with the percentage of expression of the marker in the cluster. LEC clusters are colored in blue, MEnt in green and CEnt in orange. E. Pearson correlation between pairs of ECIIa clusters. We also performed hierarchical clustering on all clusters. We outlined in red the groups of clusters showing highest correlations with one another: three out of four LEC clusters (ECIIa_0, ECIIa_1, ECIIa_6); both CEnt clusters (ECIIa_3, ECIIa_5); all MEnt clusters (ECIIa_2, ECIIa_4, ECIIa_7). One of the LEC clusters, ECIIa_8, correlated less strongly with other LEC clusters but also correlated with MEnt cluster ECIIa_4, indicating that it is a transition cluster between MEnt and LEC. The cross-correlogram is colored by Pearson correlation coefficient. The significant correlations after Benjamini-Hochberg multiple testing correction are marked with asterisks.

As a complementary analysis to the GO pathways, we also performed GSEA with WikiPathways, and found genes from the ErbB signaling pathway among the significant gene sets (padj = 0.03, NES = 1.77). Enrichment for energy production pathways emerged as well as a salient characteristic of ECIIa neurons: we found enrichment for pathways involved in electron transport chain (padj = 0.03, NES = 1.71), in glycolysis (padj = 0.09, NES = 1.68) and to a lesser extent in tricarboxylic cycle (nominal p-value = 0.03, NES = 1.51). Supporting the findings of higher expression levels for electron transport chain genes in ECIIa, we found that *Ppargc1a*, which codes for PGC-1a, a master regulator of mitochondria biogenesis (Fernandez-Marcos and Auwerx, 2011), was significantly higher in all ECIIa neurons, as well as in the subregions CEntIIa and LECIIa. In MEntIIa, the p-value of enrichment was nominally significant. Even if the effect size was modest (log2 Fold-Change of enrichment (log2FC) = 0.49, 0.52, 0.51 for LECIIa, MEntIIa, CEntIIa respectively), *Ppargc1a* transcript levels in the nucleus were consistently overexpressed in all three subregions of layer IIa. Our previous cell-type specific ribosomal profiling data showed that while *Ppargc1a* was higher in principal neurons of S1 and V1 than in ECIIa neurons, it was expressed at higher levels in ECIIa than in hippocampus (Roussarie et al., 2020). Key regulators of lipid and glucose metabolism, *Foxo1*, *Pparg* and *Hnf4g,* higher in ECIIa neurons, further supports a coordinated transcriptional signature encompassing multiple metabolic domains.

### Molecular characterization of Reln positive neurons in layer IIa

In addition to the high levels of functional specialization between MEC and LEC, different subdivisions of EC in humans also display differential vulnerability to NFT formation, with the earliest NFTs appearing in the posterior half and lateral edge of human EC (Braak and Braak, 1985, 1995; Llamas-Rodríguez et al., 2022). To gain more precise insight into the diversity of layer IIa neurons, we subclustered nuclei from final_exc2, final_exc6, and final_exc10 (*Reln*-positive excitatory neurons from LEC, MEnt and CEnt layer IIa respectively) (Fig. 3C-D). After subclustering, the distinctions between LEC, MEnt and CEnt persisted as expected but we were also able to gain much finer resolution on the different subtypes of neurons in this layer. Four of the subclusters were made of nuclei originally members of the LEC subcluster (clusters ECIIa_0, ECIIa_1, ECIIa_6 and ECIIa_8) and expressed *Erbb4* (*Erbb4* was expressed but not enriched in ECIIa_8). Two subclusters were enriched in *Lef1*, *Il1rapl2* and *Vwc2l* and corresponded to CEnt neurons (clusters ECIIa_3 and ECIIa_5). Last, nuclei from the original MEnt subcluster were found in three subclusters (clusters ECIIa_2, ECIIa_4 and ECIIa_7), two of which were enriched in *Dcn* (ECIIa_2, ECIIa_7) and two in *Adam33* (ECIIa_2, ECIIa_4). Like for all excitatory neuron clusters, we calculated pairwise Pearson correlations for whole-genome gene expression levels with each pair of ECIIa clusters. We then performed hierarchical clustering of the clusters (Fig. 3E). We found strong correlations between the three MEnt clusters, between the two CEnt clusters, and three of the four LEC clusters (although all four LEC clusters were found in the same branch in the hierarchical clustering). The correlation of ECIIa_8 with other LEC clusters was less strong, and it showed some correlation with MEnt cluster ECIIa_4.

The subclusters of ECIIa neurons could correspond to anatomical subregions within LEC, CEnt and MEnt, or alternatively, they could reflect cell states, corresponding to the past or present activity history of ECIIa neurons.

#### LECIIa subdivisions

Using ISH data from the Allen Brain Atlas, we first mined genes enriched in clusters ECIIa_0, ECIIa_1, ECIIa_6 and ECIIa_8 for genes with heterogeneous expression patterns across subregions of the LEC. For instance, we found that *Gpc5*, enriched in cluster ECIIa_0 within EC, seemed to be expressed at higher levels in dorsal subregions of the LEC. *Tenm3*, enriched in clusters ECIIa_0 and ECIIa_1 was expressed at higher levels at intermediate levels. The ECIIa neurons with highest levels of *Grik1*, expressed in all LEC, but enriched in cluster ECIIa_6, could be found in the most ventral parts of LEC. Lastly, *Stxbp6* expressing ECIIa_8 neurons were found in the most caudal parts of the LEC, where it gradually transitions into MEnt and CEnt. We therefore performed multiplexed ISH for *Gpc5*, *Tenm3*, *Grik1* and *Reln* (Supplementary Fig. 9) and obtained data from the Allen Institute ISH atlas for *Stxbp6* (Supplementary Fig. 10) on coronal sections of the mouse brain (coronal sections are best to capture the extent of the LEC, sagittal sections are preferrable for CEnt while MEnt can be visualized with both orientations. All three regions can be also adequately captured by horizontal sections). We verified the patterns seen in the Allen Brain Atlas, whereby, among Reln cells from layer IIa, the dorsal-most LECIIa neurons neighboring the rhinal fissure expressed high *Gpc5* levels but little *Tenm3*, *Tenm3* increased in intensity in the dorsomedial intermediate levels of ECII, and the ventral-most parts of ECII in the LEC expressed high levels of *Grik1*, but low levels of *Tenm3* and *Gpc5*. Parcellation of the LEC was proposed by Van Groen, with the LEC divided into three subregions: the dorsolateral entorhinal field (DLE), the dorsal intermediate entorhinal field (DIE) and the ventral intermediate entorhinal field (VIE) (Groen, 2001). The three fields were delineated based on the cytoarchitectonic properties of all layers, but layer IIa is different enough across fields to allow for their accurate identification purely based on a IIa staining. Layer IIa of DLE has sparser cells than DIE and seems more loosely organized. This was exactly the case for *Gpc5* and *Reln* positive neurons. These cells extended all the way to the rhinal fissure, including the perirhinal cortex. Layer IIa of DIE, is more densely packed, and there is “a cell-sparse area” immediately under it (Groen, 2001). This corresponded to the area of ECIIa where *Gpc5* is not expressed but *Tenm3* is expressed at high levels. Lastly, in VIE, layer IIa forms a thin continuous band of cells (Groen, 2001). This is what we observe in the sector of ECIIa that expresses high levels of *Grik1*. Overall, areas of peak expression of *Gpc5*, *Tenm3* and *Grik1* allows us to demarcate DLE, DIE and VIE respectively. We therefore confirm that these subareas that were delineated based on cytoarchitectonic features by Van Groen are good fit for the molecular subtypes of neurons in the layer IIa of LEC (with cluster ECIIa_0 corresponding to DLE cells, cluster ECIIa_1 to DIE cells and cluster 6 to VIE cells). ECIIa_8, on the other hand, might be a transition region between LEC and MEnt: it is located in the caudoventral corner of the DIE, ventral to the most caudal parts of the LEC, which we could validate in our IF staining for STXBP6 and RELN. In the initial clustering, cells from subcluster ECIIa_8 were clustering with LEC, but by decreasing the resolution of clustering, ECIIa_8 sometimes co-clusters with MEnt instead. A feature of layer II *Reln* cells in MEC vs LEC is the density and thickness of this layer. Very clearly, in *Stxbp6* negative areas from VIE, ECIIa neurons present the dense thin aspect described by Van Groen et al. whereas in the *Stxbp6* positive area, ECIIa neurons are more spaced, although grouped in patches, and the layer is thicker, akin to what Van Groen et al. have characterized as MEnt.

In the MEC, the fact that nuclei from MEnt and CEnt form several subclusters at the resolution used here indicates that both MEC subregions might be more diverse than suggested by Van Groen’s parcellation, where CEnt and Ment were not further subdivided in subareas.

#### CEntIIa subdivisions

Although Van Groen did not subdivide CEnt into subregions, the heterogeneity of the region is clear at the electrophysiological level, where studies have shown dorsoventral gradients in the properties of CEnt cells (Garden et al., 2008; Giocomo and Hasselmo, 2009). We therefore asked if our two CEnt clusters (ECIIa_3 and ECIIa_5) have an anatomical substrate. We identified two markers, *Kirrel3* and *Tenm3*, enriched in ECIIa_3 and ECIIa_5 respectively, and performed multiplexed ISH combined with IF on sagittal sections for these two genes as well as *Il1rapl2* and RELN protein to unequivocally discriminate CEntIIa from MEntIIa or LECIIa neurons (Supplementary Fig. 11). Our ISH clearly showed two distinct populations, which correspond to the two branches of the roughly L-shaped CEnt (see Supplementary Fig. 4): a *Kirrel3* positive lateral CEntIIa population (also the most ventral part of CEntIIa, the base of the L, ECIIa_3) and a *Tenm3* positive medial population (the stem of the L, ECIIa_5).

#### MEntIIa subdivisions

To identify the subdivisions of MEnt corresponding to ECIIa_2, ECIIa_4 and ECIIa_7, we analyzed again the experiment shown in Supplementary Fig. 6 (a multiplexed ISH combined with IF on coronal sections for *Cdh18*, *Dcn*, *Rfx3* and RELN protein detection) (shown in Supplementary Fig. 12). Indeed, ECIIa_4 was characterized by highest expression of *Rfx3*, while ECIIa_2 and ECIIa_7 by high expression of *Dcn*. ECIIa_7 could then be distinguished from ECIIa_2 by its expression of *Cdh18* in addition to *Dcn*. We were able to show that ECIIa_4 corresponds to the population immediately ventral to the LEC. ECIIa_2 is then located medial to ECIIa_4, and ECIIa_7 at the medial end of MEnt.

### Gradients of cell identities in ECIIa

By definition, clustering of single nuclei profiles discretizes cell population. To more rigorously assess whether the subpopulations of ECIIa neurons represent discrete cell subtypes or a continuous anatomical gradient and investigate the transitions between subareas, we performed trajectory inference analysis for all ECIIa neurons (pooling LECIIa, CEntIIa and MEntIIa nuclei). We first used the Slingshot software to search for “lineages” within ECIIa neurons in an unsupervised way. We obtained three different lineages that started from the most medial part of MEntIIa (subcluster ECIIa_7), two of them finishing in LEntIIa (subcluster ECIIa_0 or subcluster ECIIa_6 respectively), one of them finishing in CEnt (Fig. 4). We then performed gene expression modeling with tradeSeq, ordering nuclei along a pseudotime axis spanning ECIIa cells from each lineage, investigating whether gene expression changes occur gradually or abruptly along each trajectory. All transitions between clusters appeared to be gradual (Fig. 4). We hypothesized that these lineages had an anatomical substrate, and that different points of the lineage corresponded to different anatomical locations of each of the regions. We used ABC atlas data to show that this was the case (Supplementary Fig. 13-15). All three spanned subdivisions of MEnt, ECIIa_7, ECIIa_2, ECIIa_4, indeed represented progressively more lateral subdivisions of MEnt. The *MEnt* part of the lineages terminated at ECIIa_8, which as mentioned above, co-clusters with MEntIIa or with LECIIa depending on the clustering resolution. The transition from ECIIa_7 to ECIIa_8 via ECIIa_2 and ECIIa_4 appeared to be gradual, with genes with peak expression that slowly shifted across MEnt subregions (from the deep parts of MEnt that transition with layer Va (ECIIa_7), to rostro-medial parts (ECIIa_2), to lateral parts of MEnt (ECIIa_4), to caudoventral parts of LEC (ECIIa_8)). We found genes displaying expression patterns overlapping ECIIa_8 and ECIIa_4 (like *Penk*) and other genes with expression patterns overlapping ECIIa_8 and ECIIa_1 (like *Satb1*), confirming the LEC-MEnt transitional nature of cluster ECIIa_8 (Supplementary Fig. 13). The two *LEC* lineages gradually transitioned from ECIIa_8 to ECIIa_1, indicating the absence of an abrupt border between the two parts of DIE. The two lineages separated in ECIIa_1, with one lineage gradually transitioning towards ECIIa_0 (from DIE to DLE) (Supplementary Fig. 13), and the other one towards ECIIa_6 (from DIE to VIE) (Supplementary Fig. 14). The third lineage, finishing in *CEnt*, which went from ECIIa_8 to ECIIa_3 and 5 showed a smooth transition between ECIIa_8 and ECIIa_3, and another transition between ECIIa_3 and ECIIa_5 (Supplementary Fig. 15). Interestingly, within both ECIIa_3 and ECIIa_5, the trajectory analysis indicates internal gradients, that are likely to have an anatomical basis as well. Examination of the expression of genes enriched at the pseudotime start and end of each of these (*Stxbp6* and *Abi3bp* for ECIIa_3 and *Cntnap5b* and *Cpa6* for ECIIa_5) indicated that ECIIa_3, the rostroventral CEntIIa that roughly runs along a horizontal mediolateral axis presented a gradient from lateral parts (higher in *Stxbp6*) to medial parts (higher in *Abi3bp*). ECIIa_5, on the other hand, the caudomedial CEntIIa that roughly runs along a vertical dorsoventral axis, presented a gradient from dorsal (*Cntnap5b*) to ventral parts (*Cpa6*). In the most dorsal parts of CEntIIa, ECIIa_5 extends laterally as well, along the rhinal fissure; all the mediolateral extent of this dorsal section is high in *Cpa6*. Our trajectory inference suggests that ECIIa_3 and ECIIa_5 might be the two halves of a continuous gradient running from rostrolateral to dorsomedial parts of the CEnt.

**Figure 4.**
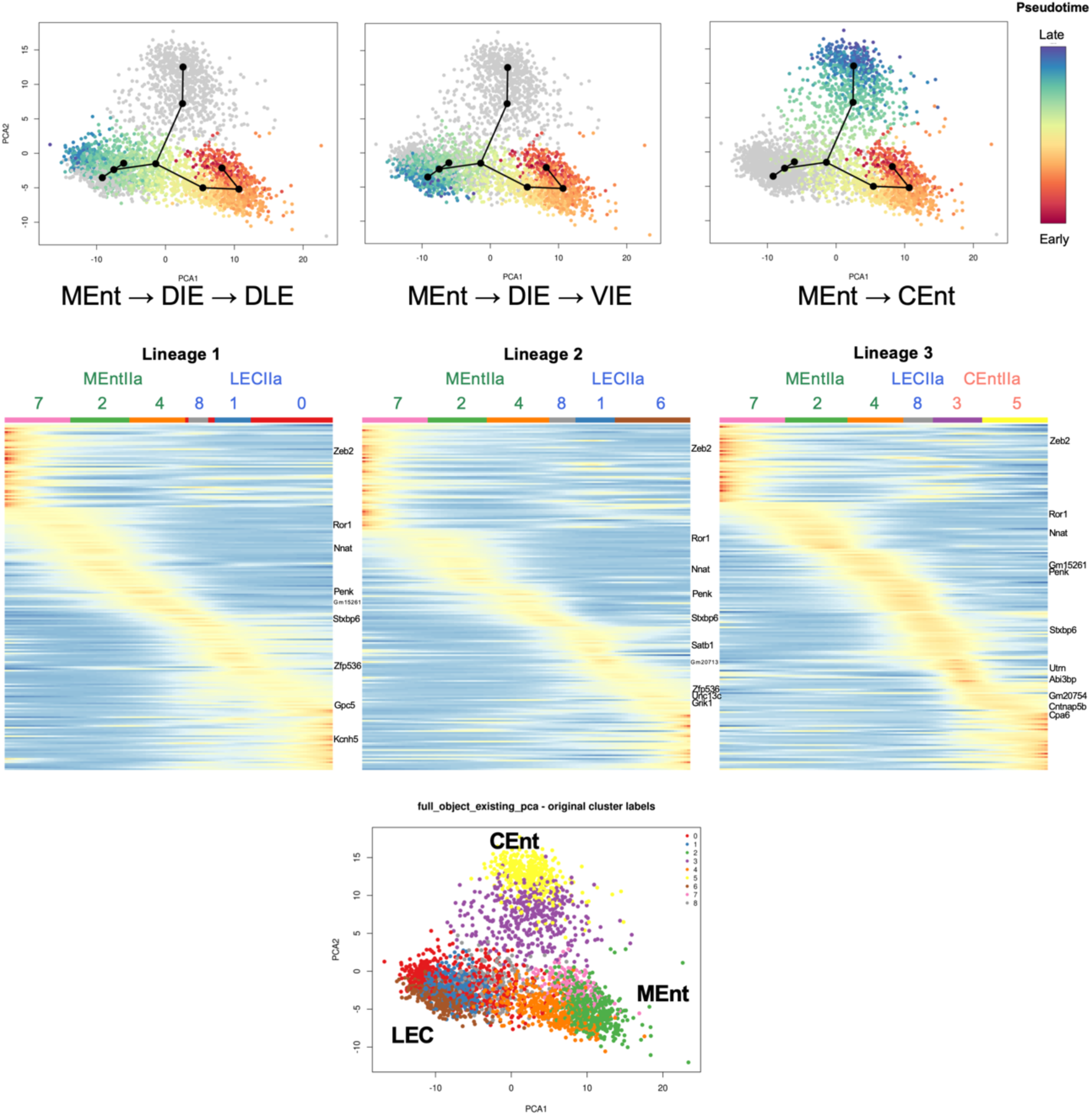
Slingshot trajectory inference analysis. We performed unsupervised trajectory inference analysis on all ECIIa neurons, using the clustering we had obtained in Figure 3 – shown at the bottom, on a PCA plot. Three different “lineages” were found, shown on the PCA plots on the top row, colored based on the pseudotime value for each cell of the lineage. The lineages were the following: lineage 1: ECIIa_7 → ECIIa_2 → ECIIa_4 → ECIIa_8 → ECIIa_1 → ECIIa_0; lineage 2: ECIIa_7 → ECIIa_2 → ECIIa_4 → ECIIa_8 → ECIIa_1 → ECIIa_6 ; lineage 3: ECIIa_7 → ECIIa_2 → ECIIa_4 → ECIIa_8 → ECIIa_3 → ECIIa_5 (or MEnt → DIE → DLE ; MEnt → DIE → VIE ; MEnt → CEnt respectively). The middle row shows the normalized expression heatmaps of the top 200 dynamic genes across each lineage. For visualization, fitted expression was z-scored independently for each gene across pseudotime. Therefore, heatmap colors represent expression relative to that gene’s own mean fitted expression rather than absolute expression differences between genes. Genes were ordered according to the pseudotime position of their maximum fitted expression, producing an early-to-late peak ordering. To relate the continuous fitted expression trajectories back to the original Seurat clusters, the heatmap columns were annotated with the dominant original cluster at each pseudotime position.

Overall, we find that ECIIa_8 is a general transitional area between LEC, MEnt and CEnt. Physically at the center between all three subregions, it has elements of the MEC (lower Reln expression, wider layer IIa, where Reln cells are more spaced) and LEC (co-clustering with LEC in our initial clustering), but does not express any of the broad markers of the subregions (no *Erbb4*, *Il1rapl2* or *Dcn*). On the LEC side, this subcluster gradually transitions from caudoventral DIE to rostral DIE and DLE on the one hand and to VIE on the other hand. On the CEnt side, it transitions into the two gradients described above. On the MEnt side, it smoothly transitions medially before it merges into the subiculum. The ECIIa thus appears to be a collection of four gradients that meet in the nexus area ECIIa_8 and more work will be needed to understand the functional implication of these subdivisions and their developmental origins.

### Pathway analysis of ECIIa subdivisions

To gain greater understanding of the molecular differences between *Reln*-positive ECIIa neurons from LEC, MEnt and Cent, we performed pairwise pseudobulk differential gene expression analysis between clusters final_exc2 (LECIIa), final_exc6 (MEntIIa) and final_exc10 (CEntIIa). We found 593 genes higher and 681 lower in LECIIa vs. MEntIIa; 356 genes higher and 332 lower in CEntIIa vs. LECIIa; 355 genes higher and 442 lower in CEntIIa vs. MEntIIa (Fig. 5 and Supplementary Table 6). We then performed GSEA with GO pathways on these comparisons. Genes from the Heparan Sulfate Biosynthetic Process pathway were significantly enriched in LECIIa (LECIIa vs. CEntIIa: padj = 0.04, NES = 2.06, LECIIa vs. MEntIIa: padj = 0.12, NES = 1.96) (with highly significant enrichment of *Ndst3*, *Ndst4*, Hs3st2, Hst3st4, Hst3st5, and Extl3 in LECII vs either subdivision). These enzymes represent many different stages of the heparan sulfate proteoglycan synthesis - addition of N-acetyl glucosamine to the GAG-CP linker by EXTL3, N-sulfation of N-acetyl glucosamine by NDST3 and NDST4, O-sulfation of the glycan chain by HS3ST2, HS3ST4, HS3ST5 (Ouidja et al., 2024). Enrichment of these genes in the LECIIa suggests a unique composition of HSPGs in these cells, and subsequently unique signaling abilities of certain diffusible ligands or extracellular matrix proteins that use HSPGs as co-receptors to increase their interaction with membrane receptors. In particular, RELN has high affinity to HSPGs which helps establish RELN gradient during development (Donato et al., 2018), and N-sulfated HSPGs like the ones formed by NDST enzymes potentiate RELN signaling (Pan et al., 2025). It is noteworthy that RELN, although enriched in all ECIIa neurons, is expressed at higher levels in LECIIa compared to MEnt (log2FC = 2.05) and CEnt (log2FC = 0.67). Together with the HSPG biosynthesis gene enrichment there, this suggests that the RELN signaling pathway might be particularly active in LECIIa neurons. For both LECIIa vs. CEntIIa and vs. MEntIIa, we found pathways linked to the dendritic compartment (albeit a different one) with a significant enrichment (postsynaptic density membrane, LECIIa vs. CEntIIa padj = 0.03, NES = 1.61; dendrite development, LECIIa vs. MEntIIa padj = 0.08, NES = 1.55). This could suggest that LECIIa neurons form higher numbers of postsynapses per neuron than in the MEC, have larger postsynaptic densities, or that there is a greater turnover of them. Subsequently performing a GSEA analysis using WikiPathways, we found a significant enrichment for genes from the Erbb pathway (LECIIa vs. CEntIIa: padj = 5.0 x 10^-3^, NES = 2.05, LECIIa vs. MEntIIa: padj = 0.09, NES = 1.85) (*Erbb4* is the most significantly enriched gene in LECIIa vs. MEntIIa and CEntIIa, and the *Erbb4* ligands *Nrg1* and *Nrg3* were both significantly enriched in LECIIa vs. the two other subdivisions). The neuregulins could function as autocrine regulators of Erbb4 activity, known to regulate excitatory synapse maturation (Li et al., 2007), and dysregulate excitability in AD (Lee et al., 2026).

**Figure 5.**
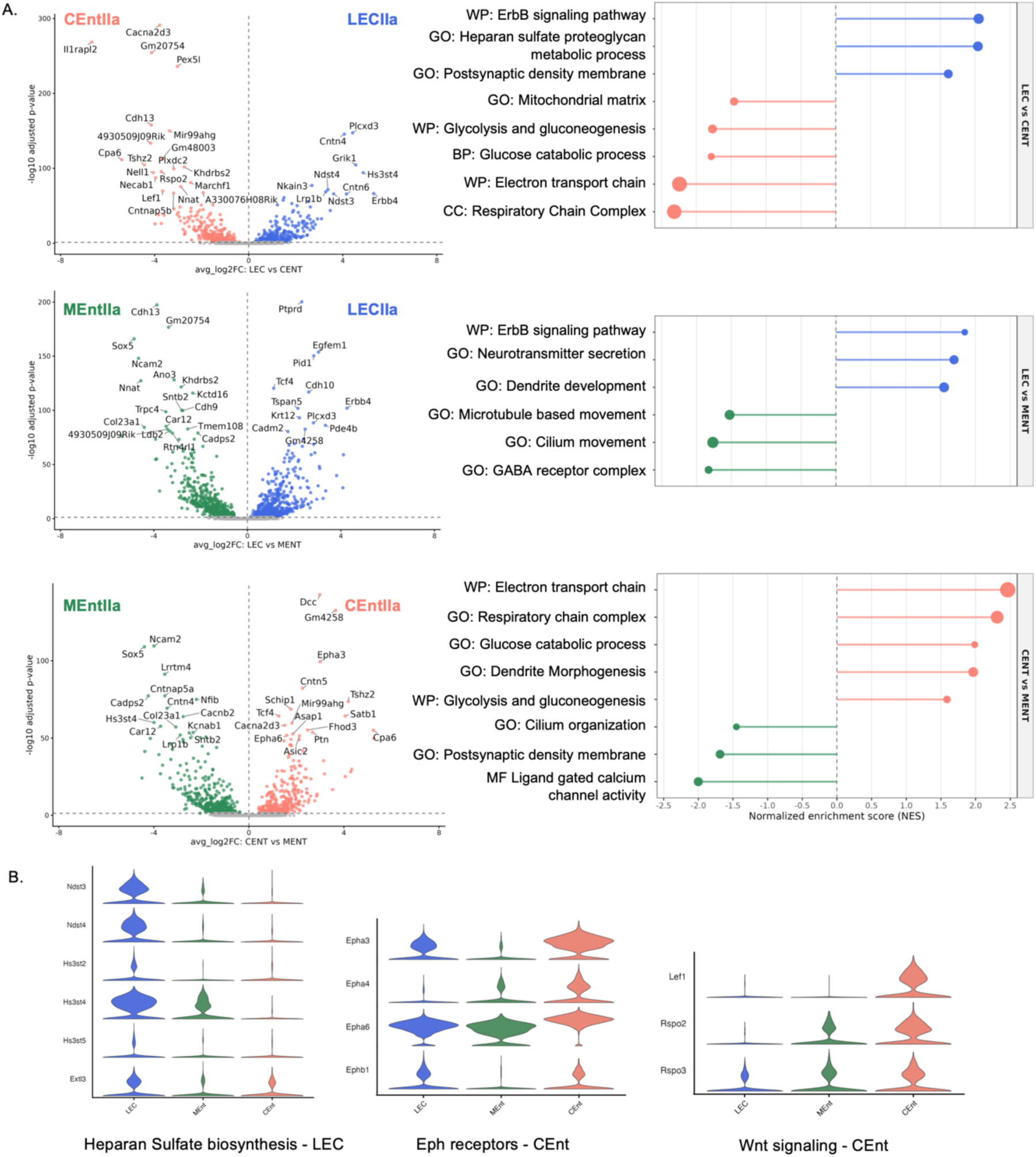
Pairwise differential gene expression analysis between LECIIa, MEntIIa, CEntIIa neurons. We pseudobulked gene expression across ECIIa_3 and ECIIa_5 for CEntIIa, across ECIIa_2, ECIIa_4 and ECIIa_7 for MEntIIa and across ECIIa_0, ECIIa_1, ECIIa_6 and ECIIa_8 for LECIIa, and used DEseq2 to find genes differentially expressed. A. Volcano plot for the differentially expressed genes for each comparison. B. Select pathways enriched in each neuron type from a GSEA analysis of the differentially expressed genes for each comparison, using GO pathways and WP. C. Violin plot for expression of heparan sulfate biosynthesis enzymes, Eph receptor genes, and Wnt signaling genes across the three cell types. Heparan sulfate biosynthesis genes are significantly enriched in LECII. We also find select Eph receptor genes, and Wnt signaling genes enriched in CEntIIa, although these latter two pathways are not significantly enriched in CEntIIa with GSEA.

Most of the pathways enriched in CEntIIa neurons were related to enrichment of various metabolic processes (e.g. respiratory chain complex, CEntIIa vs. LECIIa, padj = 1.6 x 10^-7^, NES = 2.31, CEntIIa vs. MEntIIa, padj = 2.4 x 10^-5^, NES = 2.31; glucose catabolic process, CEntIIa vs. LECIIa, padj = 0.09, NES = 1.79, CEntIIa vs. MEntIIa, padj = 0.09, NES = 1.99), suggesting that cells in CEntIIa might be more metabolically active than their LEC and MEnt counterparts. Importantly, the mice profiled in this study were housed in a conventional environment, relatively devoid of sensory stimulation. Considering the larger multisensory input in LEntIIa vs. spatial input in MEC, we might have found comparable expression of metabolic genes in mice housed in an enriched environment. Of note, a few genes central in the Wnt signaling pathway were among the top CEntIIa-enriched genes: *Lef1*, is one of the Wnt-responsive transcription factors, and *Rspo2* and *Rspo3* are two strong activators of Wnt signaling pathway. Wnt signaling, in particular the transcriptional response to Wnt has been suggested to be important for expression of activity-dependent synapse and network remodeling (Ataman et al., 2008; Gogolla et al., 2009). Wnt molecules are only detected in very sparse nuclei by snRNA-seq. On the other hand, in our bacTRAP profiling of ECIIa neurons (where we could not discriminate between LEC and CEnt) (Roussarie et al., 2020), we found some degree of enrichment of a few Wnt genes in ECIIa neurons compared to five other types of long-range projection neurons from hippocampus and neocortex (mostly *Wnt2*, log2FC = 2.4, FDR = 2.9 10^-40^; *Wnt2b*, log2FC = 0.9, FDR = 1.9 10^-12^; *Wnt5a*, log2FC = 0.8, FDR = 6.1 10^-9^; *Wnt7b*, log2FC = 0.8, log2FC = 2.6 10^-13^). Besides Wnt signaling, ephrin receptors were also among the genes most enriched in CEntIIa, with four different ephrin receptors significantly enriched there: *Epha3*, *Epha4*, *Epha6* and *Ephb1*. It is noteworthy that enrichment of both Wnt and some Ephrin receptors (*Epha3*, *Epha4* and *Epha6*) are only seen in the region (CEntIIa and not LECIIa or MEntIIa) where clear cell patches exist between Reln- and Calb1-expressing neurons, indispensable for the emergence of grid cells (Gu et al., 2018). Both signaling pathways have been found to regulate complex spatial patterns, like striatum matrix and striosomes for EphA4 (Passante et al., 2008), Ephrin-A5 for the proper patterning of the barrels in the somatosensory cortex(Prakash et al., 2000), or Wnt signaling for the development of the taste bud array on the tongue (Liu et al., 2007). If these pathways are more active during development of CEnt vs. LEC, we can speculate that they might play a role in the formation of cell patches in CEnt.

Lastly, various pathways related to cilium biology were enriched in MEntIIa, like cilium organization (MEntIIa vs. LECIIa, padj = 0.03, NES = 1.48, MEntIIa vs. CEntIIa, padj = 0.12, NES = 1.41), in line with the strong enrichment of the ciliogenesis transcription factor *Rfx3* in MEntIIa, as well as various structural cilia component like *Cfap44* or *Drc1*. In view of the concentration of GPCRs in cilia, it is possible that MEntIIa cells rely more than other ECIIa neuron on GPCR ligand signaling.

### Conservation of ECII subregion markers in primates

The EC is well conserved across mammals, as is much of the hippocampal formation. In particular, layer IIa neurons have been shown to express *Reln* in various mammalian species from ferrets(Martínez-Cerdeño et al., 2003) to beaked whales (Pérez-García et al., 2001) (as reviewed in (Kobro-Flatmoen and Witter, 2019)). Even outside the mammalian class, more distant species like salamanders have been shown to have, within the dorsomedial pallium, a layer of Reln-expressing cells with similarities with the mammalian EC(Woych et al., 2022). Beyond the mere expression of *Reln*, we sought to evaluate if the finer structure of EC was conserved between mouse and primates, in particular the subdivision between LEC and MEC. Ohara et al., by evaluating the distribution of PCP4-positive cells and telencephalon-projecting neurons in layer Vb proposed a first homology between mouse and macaque subdivisions of EC (Ohara et al., 2021). They posited that Er, Eo, Elr and the rostral part of Ei are homologues of the mouse LEC, while the caudal portion of EC, and Elc, Ec, ECl are the mouse mEC. We set out to bring additional evidence for this homology by testing expression of our ECIIa subdivision-specific markers in macaque. We performed multiplex ISH with macaque probes against markers of mouse LEC (*Erbb4*), MEnt (*Dcn*) and CEnt (*Vwc2l*) (Fig. 6). We found that these markers had well conserved patterns of expression, allowing us to confirm the previous homology between mouse and macaque subdivision based on layer V markers(Ohara et al., 2021). As shown in Fig. 6, we found that most *RELN-*positive cells from layer IIa in the most anterior sections of EC also expressed *ERBB4* – to the notable exception of the most lateral parts of EC. At intermediate antero-posterior levels of EC, and until the most posterior parts of EC, *VWC2L* was expressed in most ECIIa neurons, across the entire mediolateral span of EC. Using the subdivisions of EC first described by (Amaral et al., 1987), we found that *ERBB4* was expressed in ECIIa cells of Eo and Er, while *VWC2L* was expressed in Ei, Elc, Ec and Ecl. *ERBB4*-expressing cells of ECIIa in Eo – a thin and packed cell layer - bear striking resemblance with *Erbb4*-expressing cells of VIE in the mouse suggesting homology between the two regions. The subregion of ECIIa with DCN expression was more restricted: we found only DCN expression in the most medial part of ECIIa, the postero-medial corner of Eo with little to no co-expression of RELN, similar to mouse MEnt where Reln expression drops compared to LEC (Supplementary Fig. 16). Although it has been described in humans and not in macaques, this might be the EMI section (entorhinal medial intermediate subfield), medial to the EI subfield (Insausti et al., 1995).

**Figure 6.**
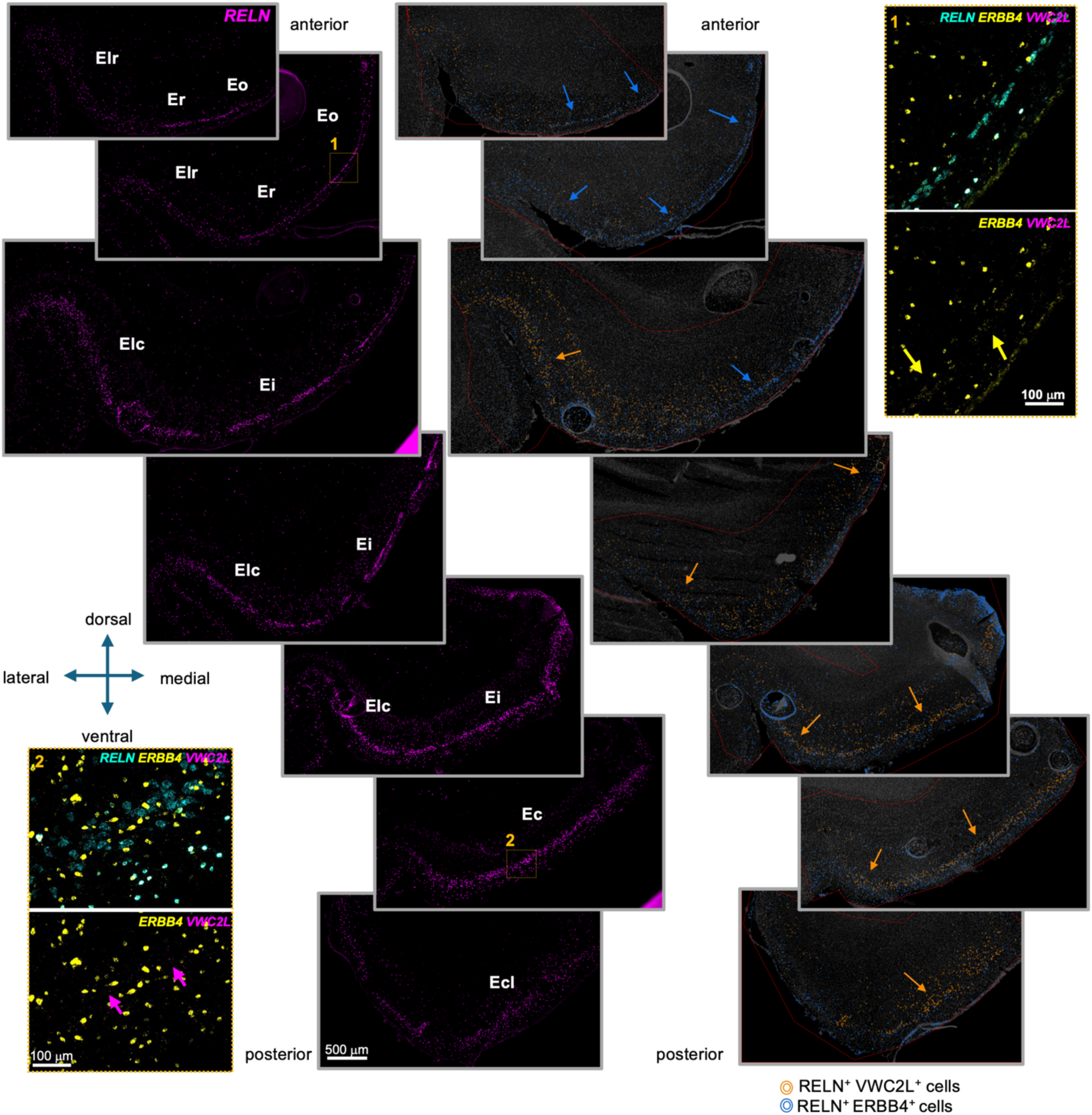
Identification of ECIIa subregions in coronal sections of 9-year-old macaque brains. ISH detection of *ERBB4*, *VWC2L*, *RELN* and *DCN*. Nuclei were counterstained with DAPI. The raw signal in two areas of ECIIa is presented on the top right of the panel (inset 1: anterior EC) and on the bottom left (inset 2: posterior EC), with RELN (cyan), ERBB4 (yellow) and VWC2L (magenta). An ERBB4+ area of layer IIa is indicated with yellow arrows. A VWC2L+ area of layer IIa is indicated with magenta arrows. B. Classifiers detecting cells expressing *RELN* and *VWC2L* (orange), and *RELN* and *ERBB4* (blue). Cells were detected using DAPI, classified based on their expression of RELN, and then further classified based on their expression of the other two markers. Coronal sections arranged from anterior-most to posterior-most and spanning the extent of the EC are shown. Images are from a male (images 1, 3, 7) and a female (images 2, 4, 5, 6) individual. Er: entorhinal rostral division, Eo: entorhinal olfactory division, ELr: entorhinal lateral division – rostral, ELc: entorhinal lateral division – caudal, Ei: entorhinal intermediate division, Ec: entorhinal caudal division, ECL: entorhinal caudal limiting division.

Additionally, we also tested if our markers with dorsoventral patterns of expression in the mouse *Gpc5*, *Tenm3* and *Grik1* presented similar expression patterns in the macaque (Fig. 7). Some differences emerged, first and foremost the absence of a notable subregion of ECIIa that would express high levels of *GRIK1* like in the mouse. The appearance of the Grik1-expression subregion (VIE) in the mouse suggests that it might be the homologue of Eo. Eo did not show any enrichment in *GRIK1* in macaque. This suggests species-specific expression in the mouse that could have significant functional consequences on the electrophysiology of mouse LECIIa cells.

**Figure 7.**
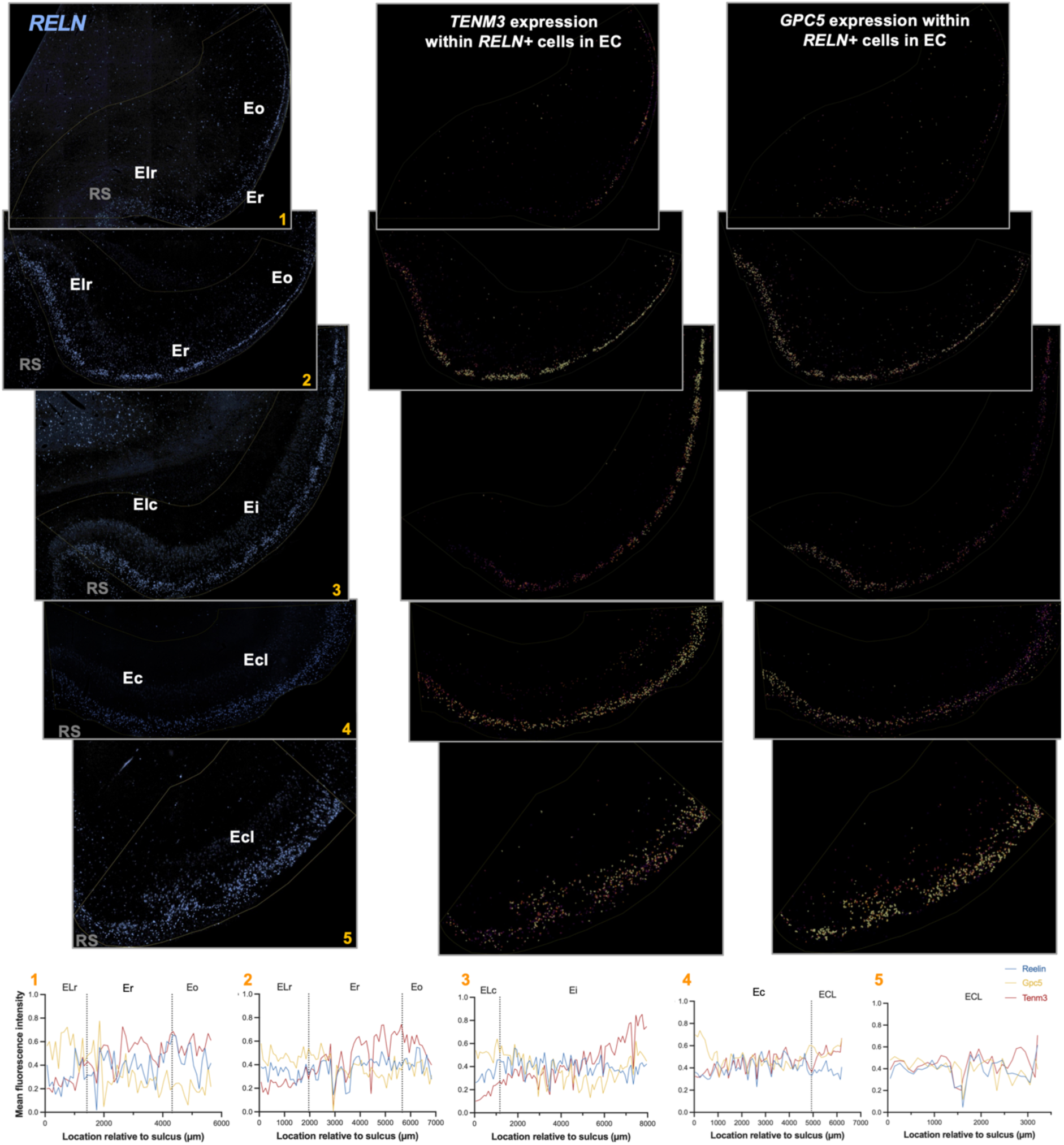
Identification of finer ECIIa subdivisions in coronal sections of a 9-year-old male macaque brain. ISH detection of *RELN*, *TENM3*, *GPC5*. Nuclei were counterstained with DAPI. The raw *RELN* signal is presented in blue on the left column. For the middle and right column, a classifier was used to detect *RELN* positive cells. Among *RELN* positive cells, we then displayed a heatmap of cell mean expression of *GPC5* (middle) and *TENM3* (left). We found that most antero-posterior levels, expression of *GPC5* and *TENM3* varied along a medio-dorsal axis. We measured, for each *RELN* positive cell, the distance to the depth of the rhinal sulcus along layer IIa. We then binned cells by segments of 100 μm along layer IIa and calculated for each bin the average cell mean expression for *RELN*, *TENM3* and *GPC5* (data shown on the bottom row). We confirmed that at most levels, there were two opposing gradients for *GPC5* on the one hand (enriched in lateral areas, closer to the rhinal sulcus) and *TENM3* and *RELN* on the other hand (enriched in medial areas, further away from the rhinal sulcus). Similar results were obtained with a 10-year-old female, a 25-year-old male and a 21-year-old female macaque brain. RS: rhinal sulcus.

On the other hand, we found very similar expression patterns for marker genes *Gpc5* and *Tenm3* in mouse and macaque EC: there were two opposing mediolateral gradients of expression for *TENM3* and *GPC5* in the most rostral parts of EC (*GPC5* highest in Elr and in the rostral part of ELc, decreasing away from the rhinal sulcus -in Er and even more in Eo; *TENM3* lowest in Elr and in the rostral part of Elc compared to Er, Eo and Ei; on the other hand, there was a relatively uniform expression of *GPC5* in caudal portions of EC like we also observed in the mouse CEnt). Altogether, our macaque data focusing on ECIIa neurons confirms the homology between LEC and rostral subdivisions of macaque Ec, and CEnt and caudal subdivisions. We also found a region with putative MEnt homology at the edge that could plausibly be the region identified as EMI in humans.

## Discussion

Here we propose a detailed molecular map of the mouse EC to start deciphering the diversity of excitatory neurons in EC, across layers, across subregions, and across anatomical territories within these subregions. The picture that emerges is one of great transcriptional heterogeneity, which is consistent with heterogeneity in various structural and functional levels (morphological, electrophysiological, cytoarchitectonic, disease vulnerability, etc…). More work is needed to grasp the significance of this molecular heterogeneity, but it underscores the value of working with precisely defined subregions for experiments involving the EC. While making the distinction between MEC and LEC is common practice (and necessary since we found that layer IIa cells in CEnt and LEC differ by hundreds of differentially expressed genes), a precise mapping within LEC for instance is rarely done. Whether in mice, macaque or human, experiments would benefit from analyzing consistent levels across the very heterogeneous EC and from using subregion specific markers. We provide markers for cell subtype identification, between and within these areas, that can be used in multiplex ISH.

We find various levels of excitatory neuron type diversity across subregions in different layers. For instance, layers Va or VI, at the resolution we consider, did not differ between regions and Vb was very similar in LEC and CEnt. In superficial layers, on the other hand, we find that the area is best subdivided into three subdivisions, namely the LEC, the medioventral EC and the caudomedial EC. The medioventral EC appears as such in the Allen reference atlas but is sometimes lumped with the caudomedial EC into the MEC. Our transcriptomic evidence shows that the two subregions are distinct, more so than subdivisions of LEC or CEnt are compared to one another. Future research will be required to determine if the medioventral section of the EC shares with CEnt some of its electrophysiological, morphological and connectivity characteristics. One major distinction between LEC and MEC neurons from layer IIa in the mouse is their terminal zone, with axons from LECIIa forming synapses onto dentate gyrus granule cells in the outer molecular layer vs. middle molecular layer for axons from MECIIa. A molecular interaction has been shown to underlie the segregation: the Netrin-G / Netrin-G ligand (NGL) interaction(Nishimura-Akiyoshi et al., 2007). *Ntng1* and *Ntng2*, enriched in LECIIa and CEntIIa respectively (as we verify in our dataset), interact with NGL-1 and NGL-2 which are localized in the outer and middle molecular layer of DG, in granule cell dendrites. Interestingly, MEntIIa expresses both Ntng1 and Ntng2. One can speculate that the projections originating from there might not be as well segregated between outer and middle molecular layers of DG as the ones from LECIIa and CEntIIa.

Honing in on cell heterogeneity within subareas, we find that previous subdivisions of the LEC fit relatively well with transcriptional patterns. Indeed, in ECIIa, the cytoarchitectonically defined DLE, DIE, and VIE match transcriptional clusters, except for a transition region between all three subdivisions of EC (cluster ECIIa_8), distinct from the rest of LEC and that is part of DIE in the Van Groen segmentation. On the other hand, we find that the CEnt subdivision is more complex than initially thought. A limitation of our dataset is that it misses neurons from the most dorsal parts of CEnt due to the dissection scheme used to isolate the EC. As a result, although we find some gradient between areas expressing high *Cntnap5b* and high *Cpa6*, we do not fully capture the entire extent of the dorsoventral MEC gradient. We find another gradient, oriented along the mediolateral axis in the most ventral sections of CEnt, which we think is separate from the usual dorsoventral gradient. It clusters separately from it and expresses a number of distinct markers. Profiling more cells from the entire extent of CEnt would allow the determination of whether there is one long continuous gradient or if the structure of CEnt is more intricate, akin to that of LEC.

Little is known about the transcriptional homology between mouse and primate EC. Architectonic studies have suggested that the primate EC is much more complex than the rodent EC, with 5 proposed subdivisions by Van Groen et al, while Amaral proposes 7 subregions – or 8 as proposed by Ohara et al who found evidence that one of the 7 regions should be split into 2 (Ohara et al., 2021). While integration of primate and mouse datasets with comparative genomics tools will be necessary to determine the full level of molecular homology between subregions of EC, our data suggest 1) more complexity than previously appreciated in the mouse and 2) high levels of transcriptional conservation between mouse and macaque. We find that markers differentially expressed in mouse LEC vs MEnt vs CEnt are also differentially expressed between EC subareas in macaques. Using layer V markers, Ohara et al identified the macaque homologues of mouse LEC and MEC. Using layer IIa markers, our data agrees with the Ohara data: with the rostral subdivisions of EC (Elr, Er, Eo and the rostral part of Ei) being the homologs of LEC, while the caudal subdivisions of EC (caudal part of Ei, Elc, Ec, Ecl) would be the homologs of CEnt. Expression of DCN in layer II at the medial edge of ECIIa suggests that there might also be a primate homologue of the medioventral section of EC, potentially the region described by Insausti as EMI (medial intermediate subfield)(Insausti et al., 1995). Study of the expression of additional markers of Ment will be needed to fully delineate its extent. Furthermore, we found very similar expression patterns for marker genes *Gpc5* and *Tenm3* in mouse and macaque EC, with opposing mediolateral gradients of expression for *TENM3* and *GPC5* in the most anterior parts of EC (*GPC5* highest in Elr and decreasing away from the rhinal sulcus, in Er and even more in Eo; *TENM3* lowest in Elr and Elc compared to Er, Eo and Ei; more uniform expression of *GPC5* in caudal portions of Ec). In the mouse, using trajectory inference analysis we provided data arguing for gradients rather than strict borders between regions.

In the macaque, quantifying TENM3 and GPC5, we also find smooth gene expression gradients for the two genes. This suggests that discrete subregions, while helpful to situate a section within the entorhinal area, misrepresent the real transcriptional nature of the area. At least in layer IIa, these gradients are organized along axes that run parallel (determining the CEnt vs. LEC expression) and perpendicular to the rhinal fissure (that distinguishes the DLE vs. DIE).

Understanding the molecular diversity of neurons within a specific brain area is not only important for cell type identification and taxonomy. It is also the first step for uncovering the molecular underpinnings of functional differences between the different cell types. Focusing on ECIIa neurons, we found key differences with the potential to determine function. Interestingly, the marker of all ECIIa neurons, Reln, with 10.9-fold higher mRNA levels in the nucleus of ECIIa neurons compared to other excitatory neurons of the area, is also enriched in LECIIa vs. CEntIIa neurons (1.6 fold more in LECIIa than in CEntIIa, 4.1 fold more in LECII than in MEntIIa). Enrichment is conspicuous at the protein level as well and could equate with higher levels of Reln signaling in this area. Interestingly, in humans, it is in posterior parts of EC, homologs of the MEC, that NFTs are first accumulating. Reelin signaling is thought to dampen tau phosphorylation (Hiesberger et al., 1999; Beffert et al., 2002; Ohkubo et al., 2003), so posterior EC might be inherently less protected against tau hyperphosphorylation, and this region would be the first to form NFT when RELN signaling starts to decrease in AD (Rodriguez-Rodriguez et al., 2025). Perhaps relatedly, we found enrichment for genes involved in heparan sulfate proteoglycan biosynthesis in LECIIa. Since many of the corresponding enzymes are enzymes that modify the particular nature of HSPGs rather than enzymes that permit their synthesis, it is likely that the flavor of HSPGs more than their abundance is different in the LEC. They function as co-receptors for a variety of diffusible ligands so this might inherently change the sensitivity of ECIIa neurons to growth factors or guidance molecules. RELN displays high affinity to HSPGs (Donato et al., 2018), so this might also contribute to higher activity of the RELN pathway in LECIIa neurons.

Beyond *Reln*, key genes are differentially expressed between ECIIa neurons of different EC subregions: expression of *Grik1*, the gene encoding the GluK1 receptor, formerly known as GluR5, has been shown to decrease synaptic vesicle release probability when expressed presynaptically (Vignes et al., 1998). We can speculate that expression of GluK1 in the lateral perforant path could be responsible for the long known difference between paired-pulse facilitation in lateral perforant path vs. long-term depression seen in medial perforant path in the dentate gyrus(McNaughton, 1980). *Ano3*, recently found to encode a neuronal activity-induced eat-me signal(Shroff et al., 2026) is highly impoverished in LECIIa neurons compared to other ECIIa neurons, to other EC excitatory neurons, and in fact also compared to long-range excitatory neurons of other brain regions that we had profiled with cell-type specific ribosomal profiling (Roussarie et al., 2020). One can speculate that this could permit long-term persistence of synapses in the area and help store stable associative memories between sensory modalities. A long-known difference between IIa neurons of MEC and LEC is the existence of a larger HCN1-mediated sag in MEC than in LEC (Nolan et al., 2007; Canto and Witter, 2012b, 2012a). This region-specific difference is likely due to significantly higher expression of *Hcn1* in CEntIIa neurons (1.6-fold), but also likely regulated by TRIP8B (encoded by *Pex5l*, even more highly enriched (8.2-fold) in CEnt vs. LEC cells) that was shown to gate HCN channels in a splice isoform dependent manner(Santoro et al., 2009). Relatedly, posttranscriptional regulation of gene expression, not captured here will add another layer of functional complexity to neuronal subtypes. We find highly differentially expressed splice factors in CEntIIa and LECIIa neurons (*Khdrbs2*, *Rbfox1* enriched in CEntIIa) as well as miRNA host genes (*MiR99ahg*, *Gm4258* also enriched in CEntIIa). Other approaches than nuclear RNA profiling, like cell-type specific ribosomal profiling, will have to decipher how these factors shape the molecular make-up of ECIIa neurons in a subregion specific manner.

Lastly, we find transcriptional evidence for higher metabolic activity of neurons from ECIIa compared to other excitatory neurons of the region. Multiple levels of poststranscriptional regulation govern its activity(Fernandez-Marcos and Auwerx, 2011), but the gene coding for PGC-1a is significantly higher expressed at the RNA level in ECIIa neurons. Overexpression of *Ppargc1a* and of bioenergetic pathways mentioned above are consistent with the enrichment of mitochondrial genes found in ISH by (Ramsden et al., 2015), and with the high levels of cytochrome oxidase levels and activity seen in layer II in rodents (Hevner and Wong-Riley, 1992; Burgalossi et al., 2011), a conserved feature in monkeys and humans (Hevner and Wong-Riley, 1992). The EC also had the highest mitochondria tissue density compared to other brain regions in a survey of mitochondria across the human brain (Mosharov et al., 2025). In our cell-type specific ribosomal profiling of ECIIa neurons, we had found several genes related to mitochondria function that were higher in ECIIa – the helicase PIF1 involved in mitochondrial DNA maintenance (Bannwarth et al., 2016) and the regulators of mitophagy OPTN(Heo et al., 2015; Tsagkogianni et al., 2026), PPEF2 (Wall et al., 2019) and TXNIP(Su et al., 2020) (cf. alz.princeton.edu). *Optn* and *Pif1* are enriched in ECIIa neurons in snRNAseq, *Ppef2* and *Txnip* are not, and their enrichment in the cell-type specific ribosomal profiling data might indicate that they are instead more translated in ECIIa than in other cells. But overall, enrichment of these genes suggests greater mitochondria content, higher mitochondria turnover, and higher metabolic needs than in other neurons. More work is needed to understand what drives these needs, whether the size of the ECIIa axonal arbor – a structural property shared with dopaminergic neurons and that people have linked with the vulnerability of these neurons to Parkinson’s disease(Pacelli et al., 2015); the high plasticity needs of these neurons, linked to their role in new memory formation, or the energy needed to make high levels of the 400kDa highly glycosylated RELN protein. Whatever its cause, high energy production comes at a cost, with potentially higher reactive oxygen species generated as a by-product, which might accelerate pathological changes in Alzheimer’s disease and contribute to the high vulnerability of this neuronal population.

## Table description

Cell type annotations (tab 1) and enriched marker genes (tab 2) for each cluster for the final clustering of excitatory neurons of EC. Differentially expressed genes between ECIIa neurons and other excitatory neurons (tab 3). Cell type annotations (tab 4) and enriched marker genes (tab 5) for each cluster for the for the ECIIa clustering. Differentially expressed genes in the pairwise comparison between ECIIa regional subtypes (tabs 6-8).

## Supplementary Figure Legends

**Supplementary Figure 1.**
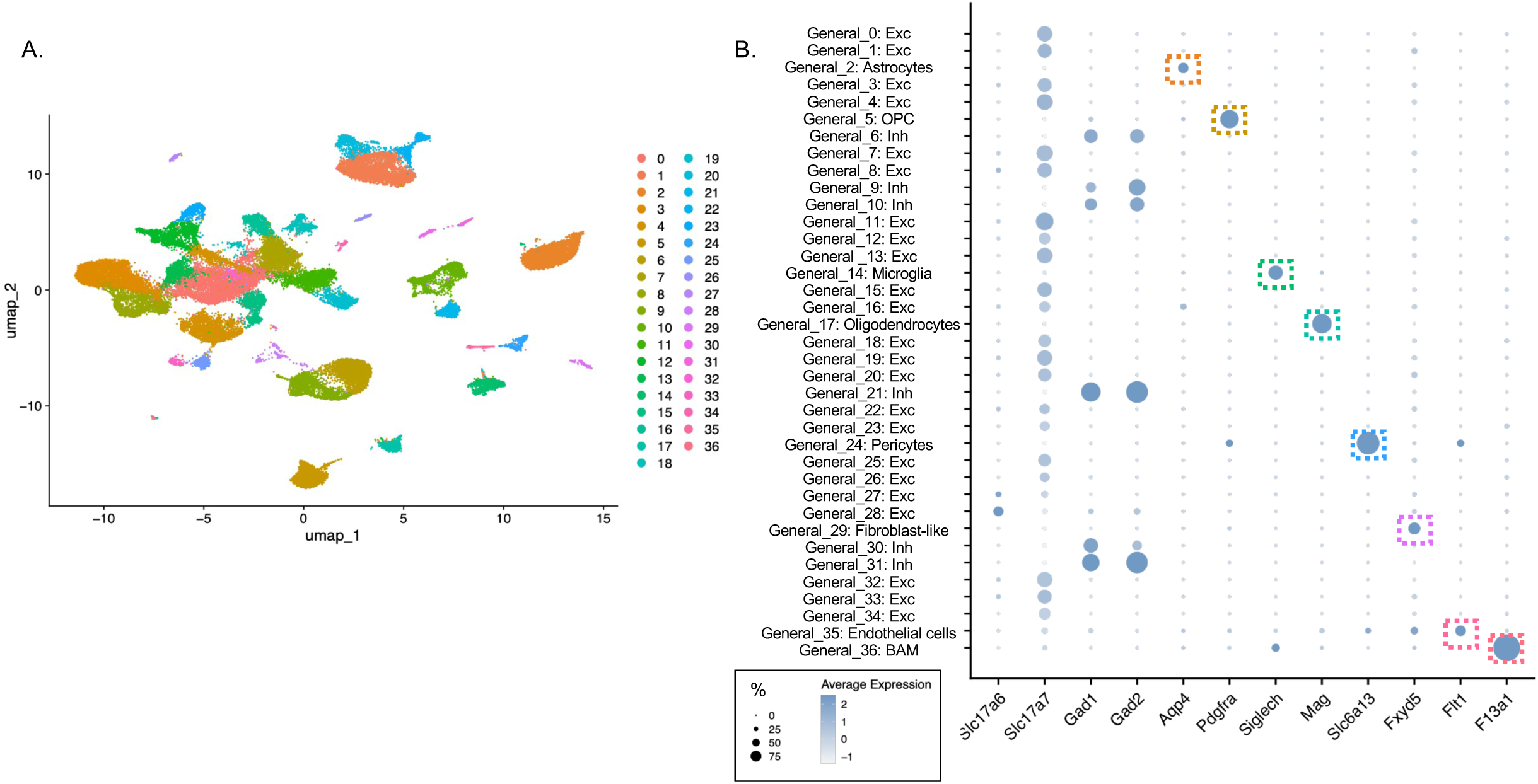
Clustering of all EC cells. We performed a general clustering of all cells that passed quality check. A. UMAP visualization of the 37 different cell clusters. B. Dot plots for general cell type specific markers in each of the 37 general clusters. Dots are colored by level of expression of the marker in the cluster, and their size correlates with the percentage of expression of the marker in the cluster.

**Supplementary Figure 2.**
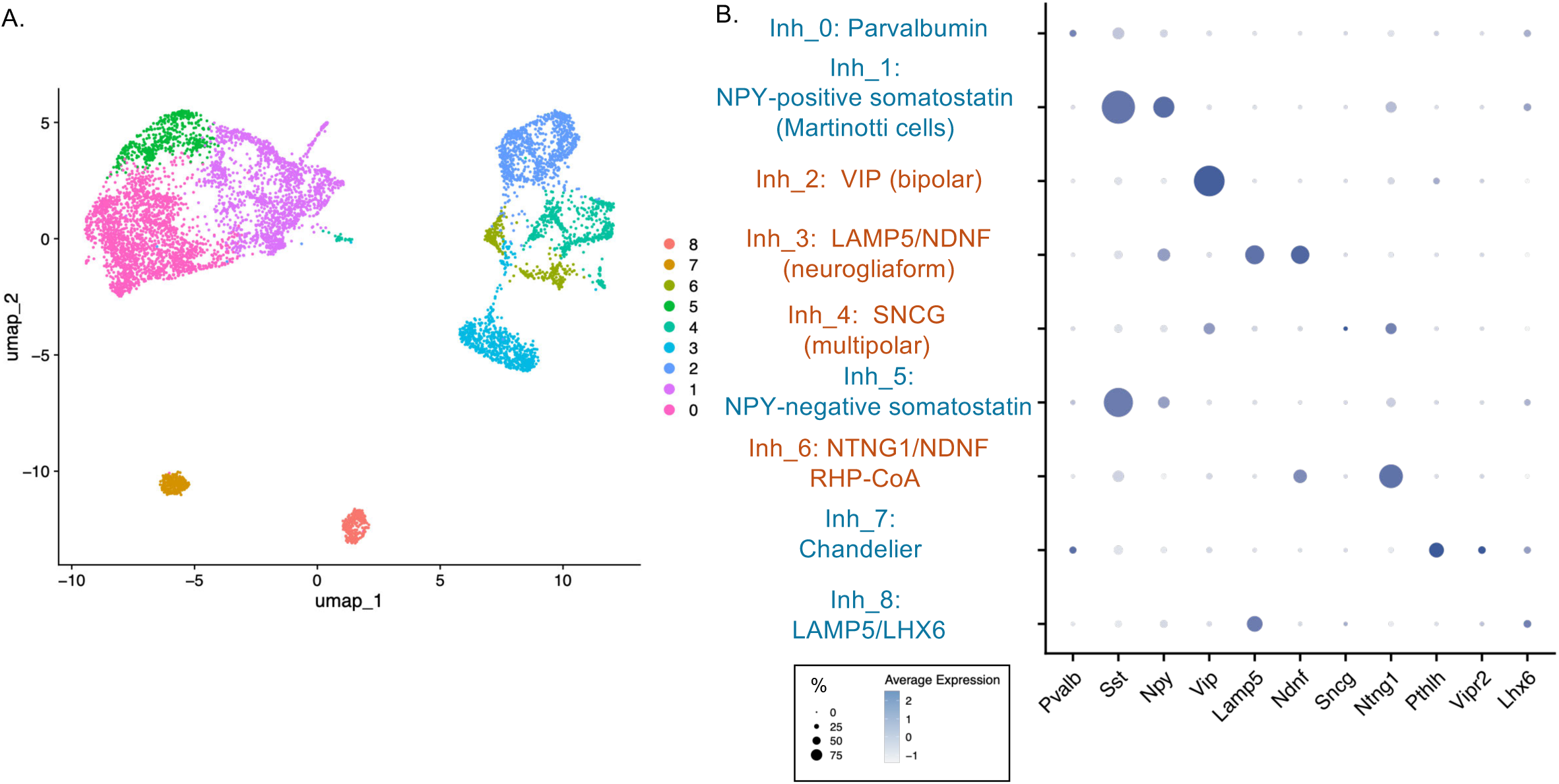
Clustering of all inhibitory neurons. We performed a reclustering of all cells with expression of Gad1 and/or Gad2 (inhibitory neurons) identified in the general clustering. A. UMAP visualization of the 9 different cell clusters. B. Dot plots for general inhibitory neuron type specific markers. Dots are colored by level of expression of the marker in the cluster, and their size correlates with the percentage of expression of the marker in the cluster. We colored medial ganglionic eminence-derived inhibitory neurons in blue, and caudal ganglionic eminence-derived inhibitory neurons in red.

**Supplementary Figure 3.**
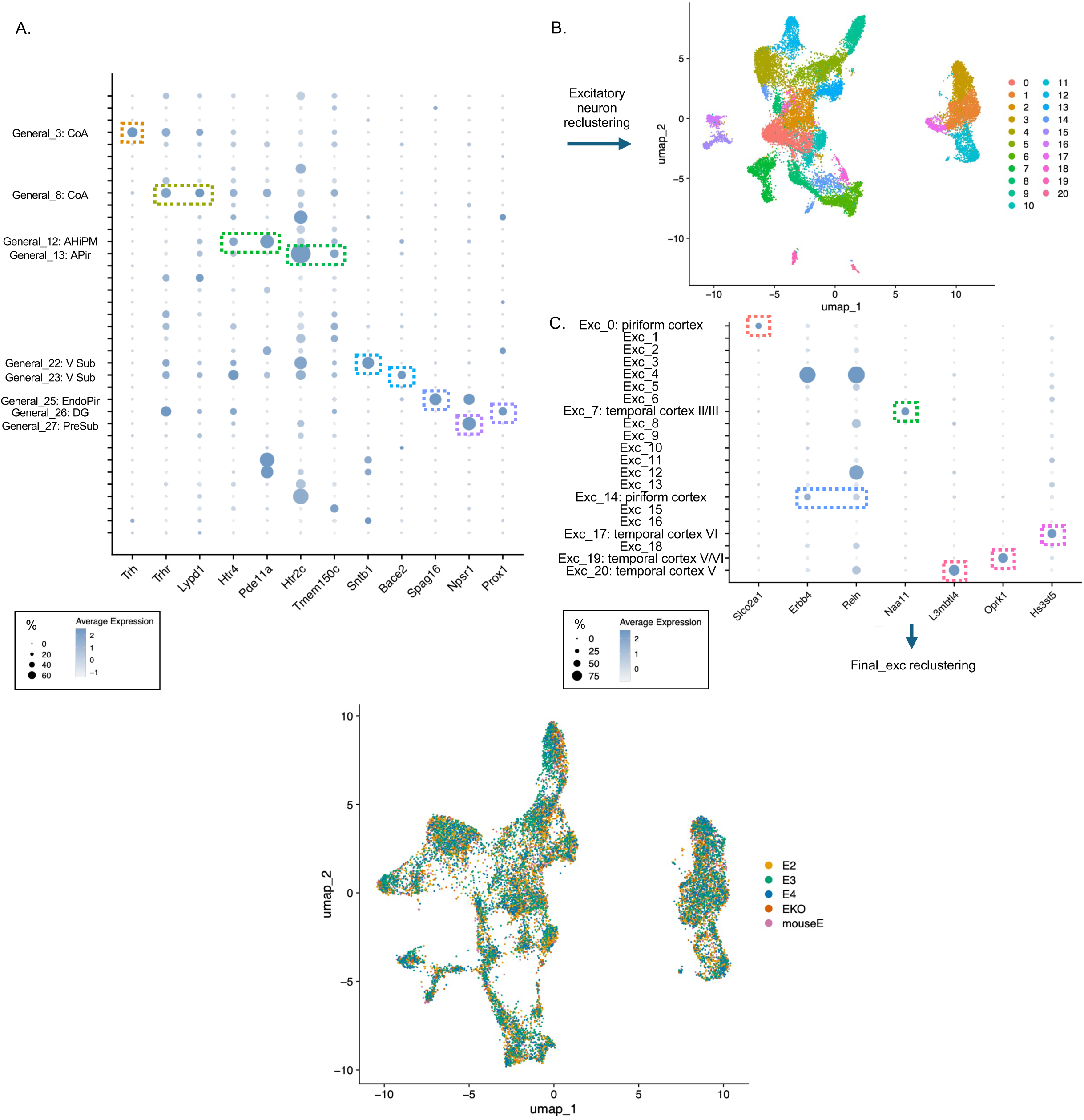
Filtering of non-EC clusters. A. Dot plots for non-EC excitatory neuron clusters in the general clustering. We show expression of marker genes for these non EC populations. All populations outlined with a dashed line were filtered out for the excitatory neuron clustering. B. UMAP for the reclustering of excitatory neurons. C. Dot plots for markers of residual non EC excitatory neuron population. Again, the clusters that are outline with a dashed line were filtered out in the final excitatory neuron clustering.

**Supplementary Figure 4.**
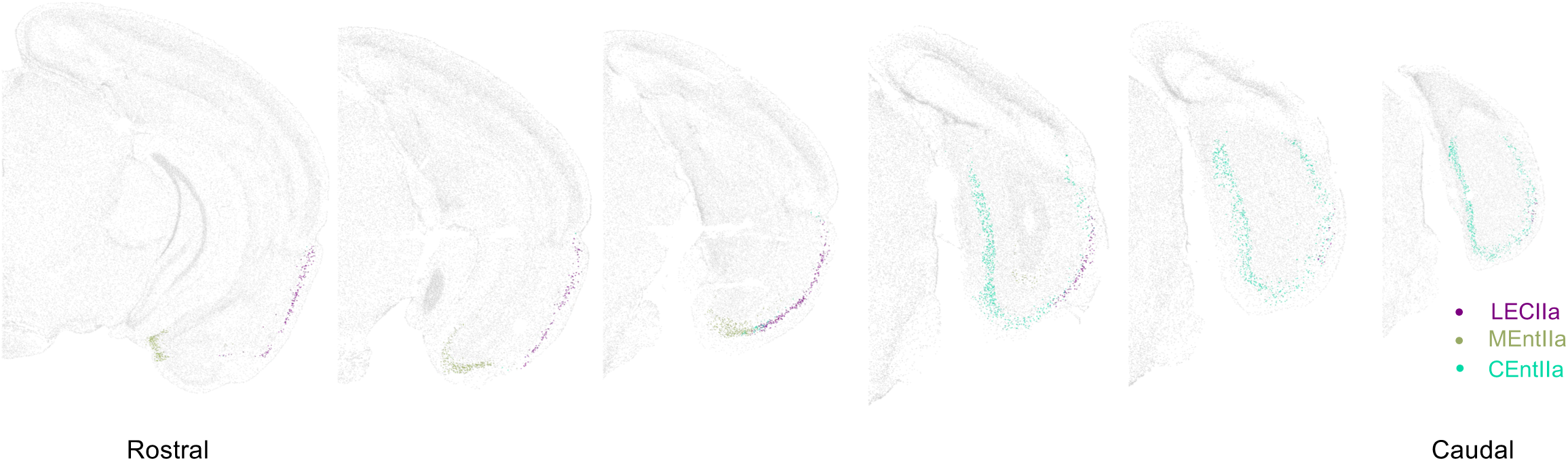
Location of cell supertypes corresponding to our LECIIa, CEntIIa, MEntIIa in the Allen ABC atlas. Using the Allen online tool, we highlighted in purple LECIIa neurons (supertype 0042), in green MEntIIa (supertypes 0066, 0067, 0068) and in cyan CEntIIa (supertypes 0051, 0052, 0053, 0054).

**Supplementary Figure 5.**
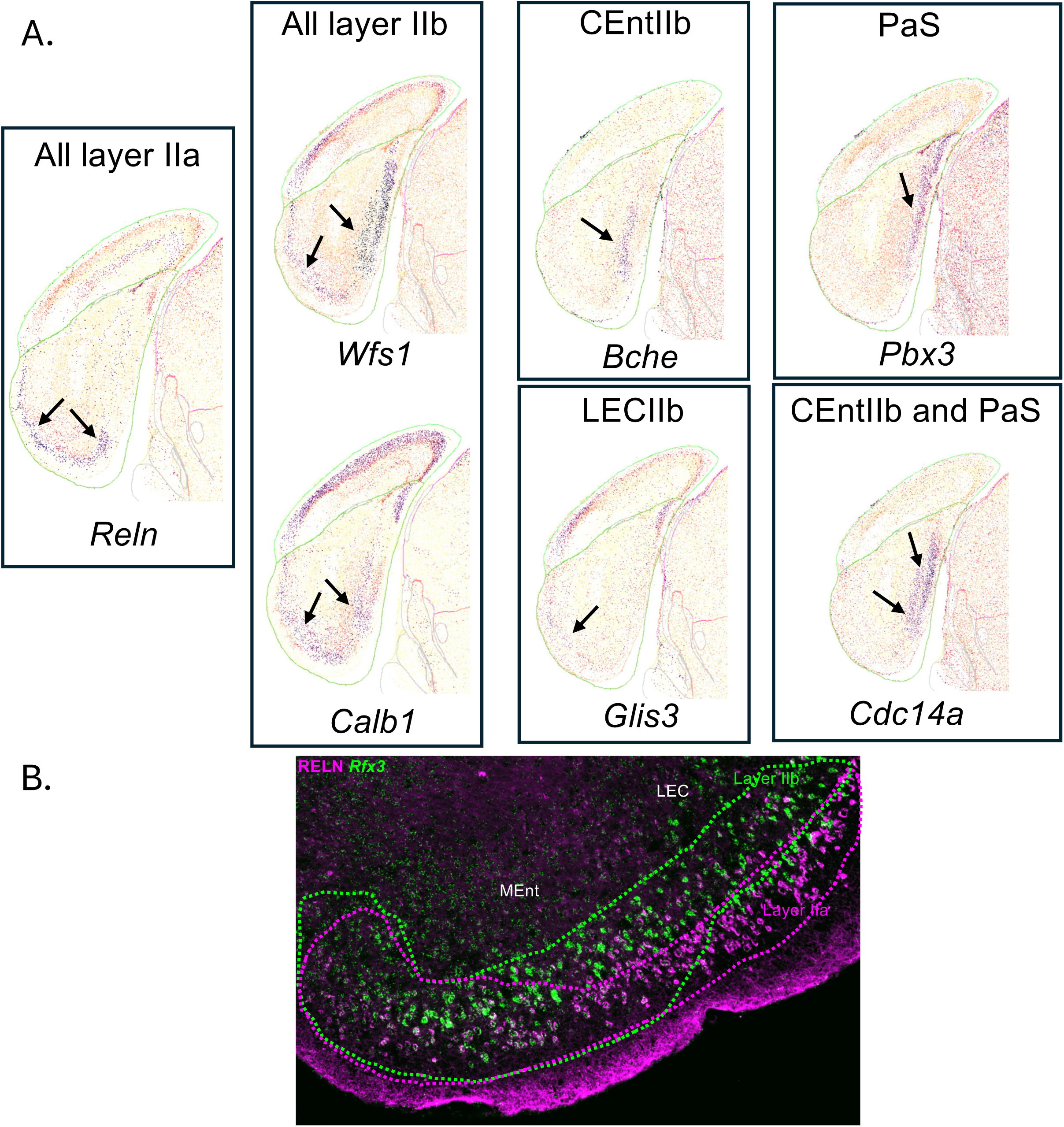
Location of cell subtypes from EC layer II. A. Heatmap images from the Allen ABC spatial transcriptomics with imputed expression atlas, showing expression level of markers of various layer II cell population: *Reln* (all layer IIa), *Wfs1* and *Calb1* (all layer IIb), *Bche* (CEntIIb or islands), *Glis3* (LECIIb), *Pbx3* (parasubiculum or PaS), *Cdc14* (CEntIIb and PaS). All images are taken from the same section of the atlas. B. Co-detection of RELN in IF (magenta) and *Rfx3* (green) in ISH in a coronal section of a 3-month old C57Bl/6J female mouse brain. In LECIIb, *Rfx3*-positive cells are found in a distinct layer from RELN cells in LECIIa, immediately beneath them. In MEntIIa, a thick layer of cells seem to co-express both RELN and *Rfx3*. At the border, the two layers seem to merge into one.

**Supplementary Figure 6.**
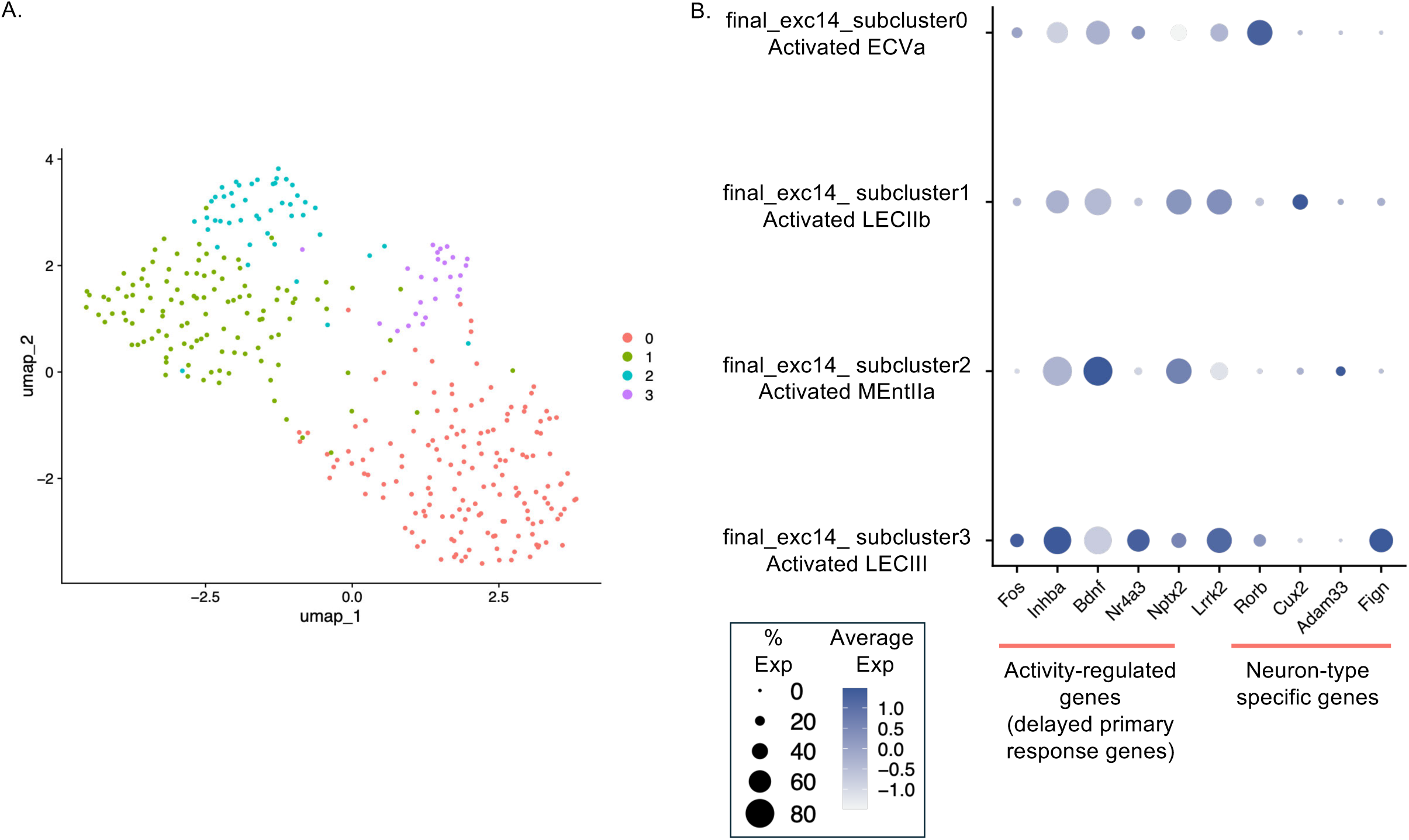
Reclustering of final_exc14. We reclustered final_exc14, a cluster enriched in activity-induced genes. A. UMAP of the 4 clusters identified after reclustering. B. Dot Plot showing that these 4 clusters are 4 distinct subpopulation of EC excitatory neurons that all express similar neuronal activity induced genes, specifically delayed primary response genes.

**Supplementary Figure 7.**
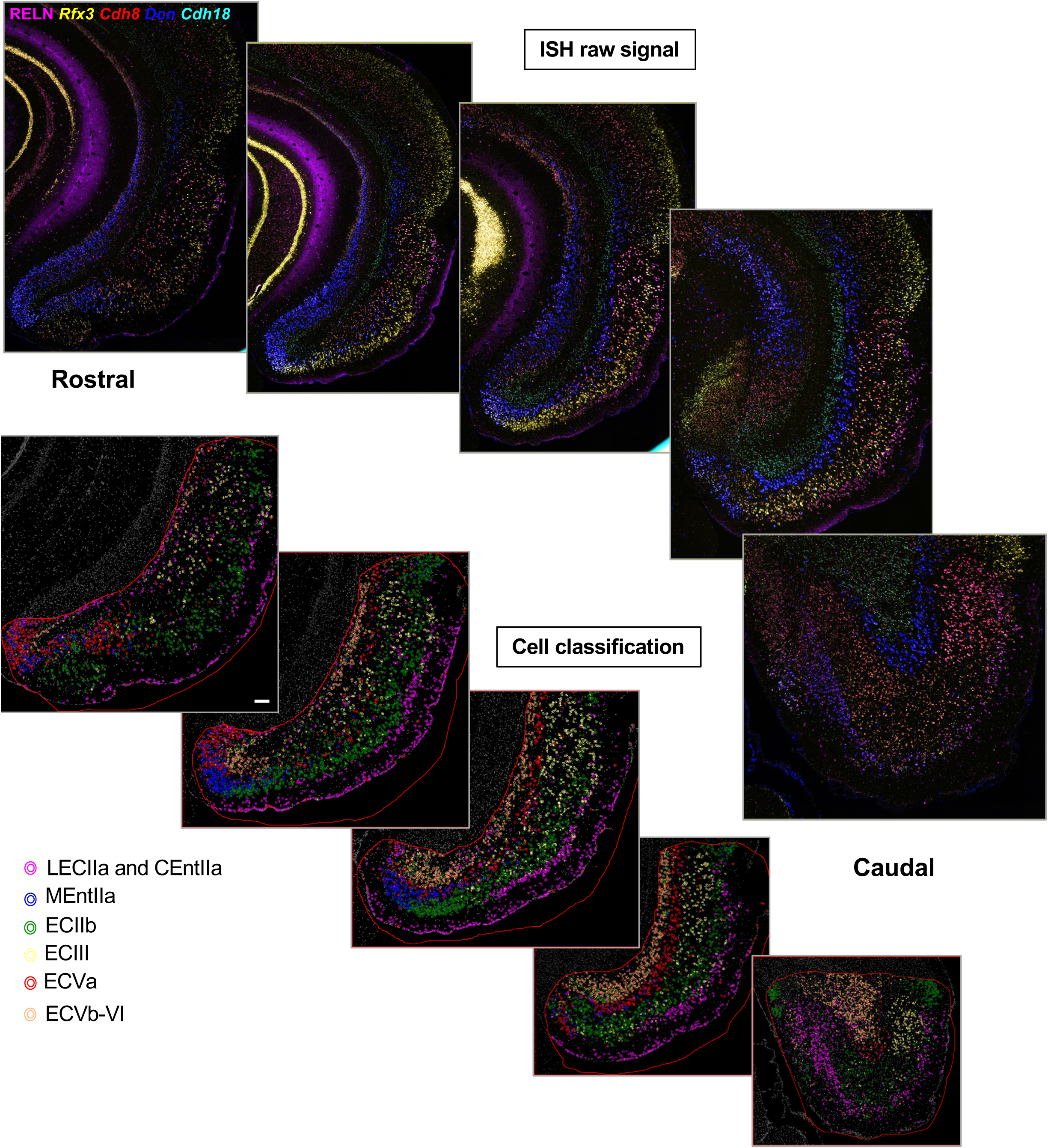
Identification of the main layers of EC in coronal sections of a 3-month old C57Bl/6J female mouse brain. A. Co-detection of RELN in IF and Rfx3 (yellow), Cdh8 (red), Dcn (blue), Cdh18 (cyan) in ISH at various rostrocaudal levels of EC (presented from rostral-most to caudal-most levels). Nuclei were counterstained with DAPI. B. Classifiers detecting cells expressing RELN (LECIIa and CEntIIa, magenta), RELN and Rfx3 (MEntIIa, blue), Rfx3 with no RELN (ECIIb, green), Cdh8 (ECIII, yellow), Dcn (ECVa, red), Cdh18 (ECVb-VI, peach).

**Supplementary Figure 8.**
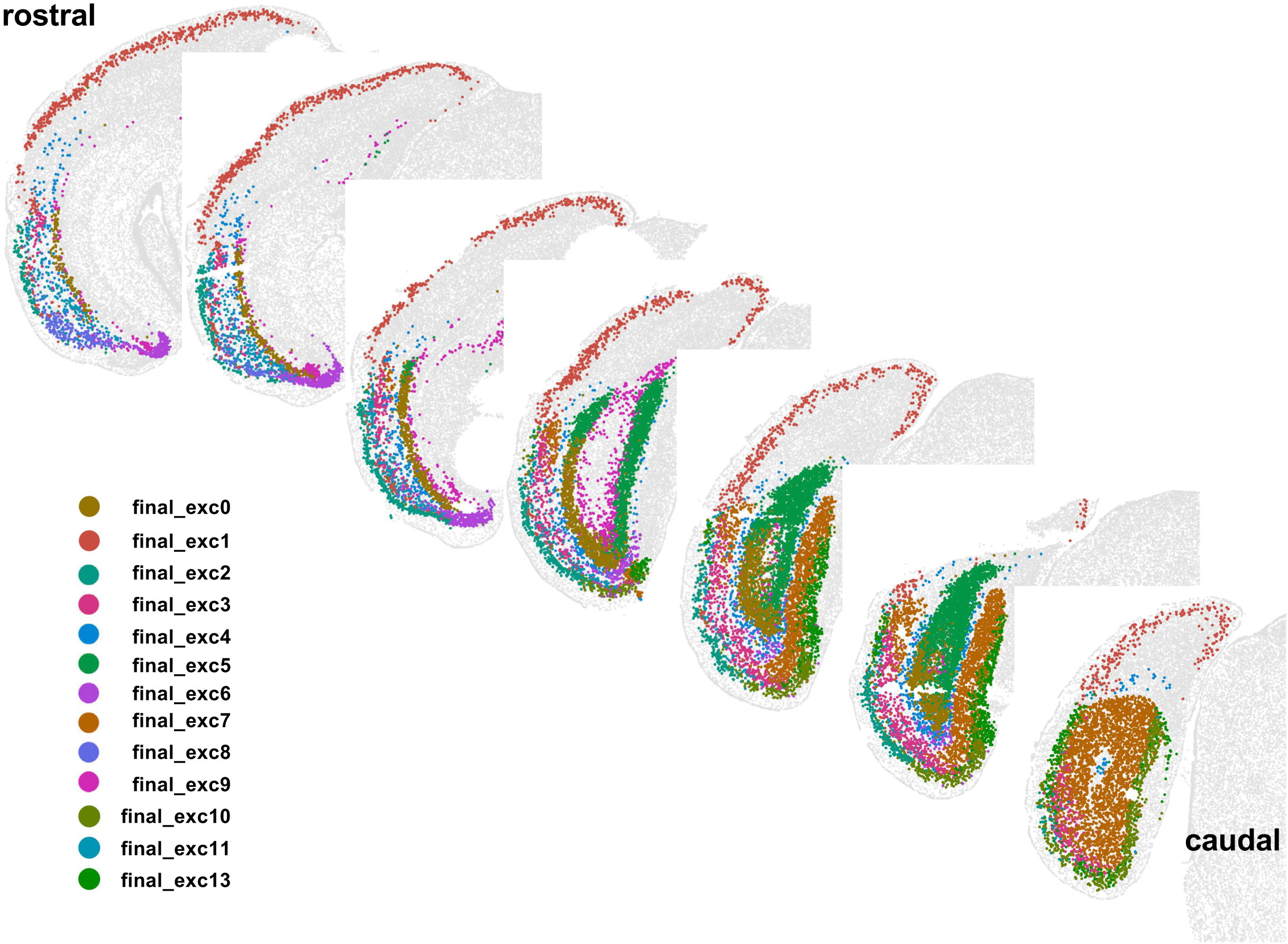
Mapping onto coronal sections from the Allen ABC atlas of our clusters. We used the Allen MapMyCells tool to identify the Allen supertype label for each of our cells. We identified the Allen supertypes represented in all of our clusters, and represented all of the supertypes corresponding to each cluster in different colors, to visualize at different rostrocaudal levels the putative location of each of our clusters.

**Supplementary Figure 9.**
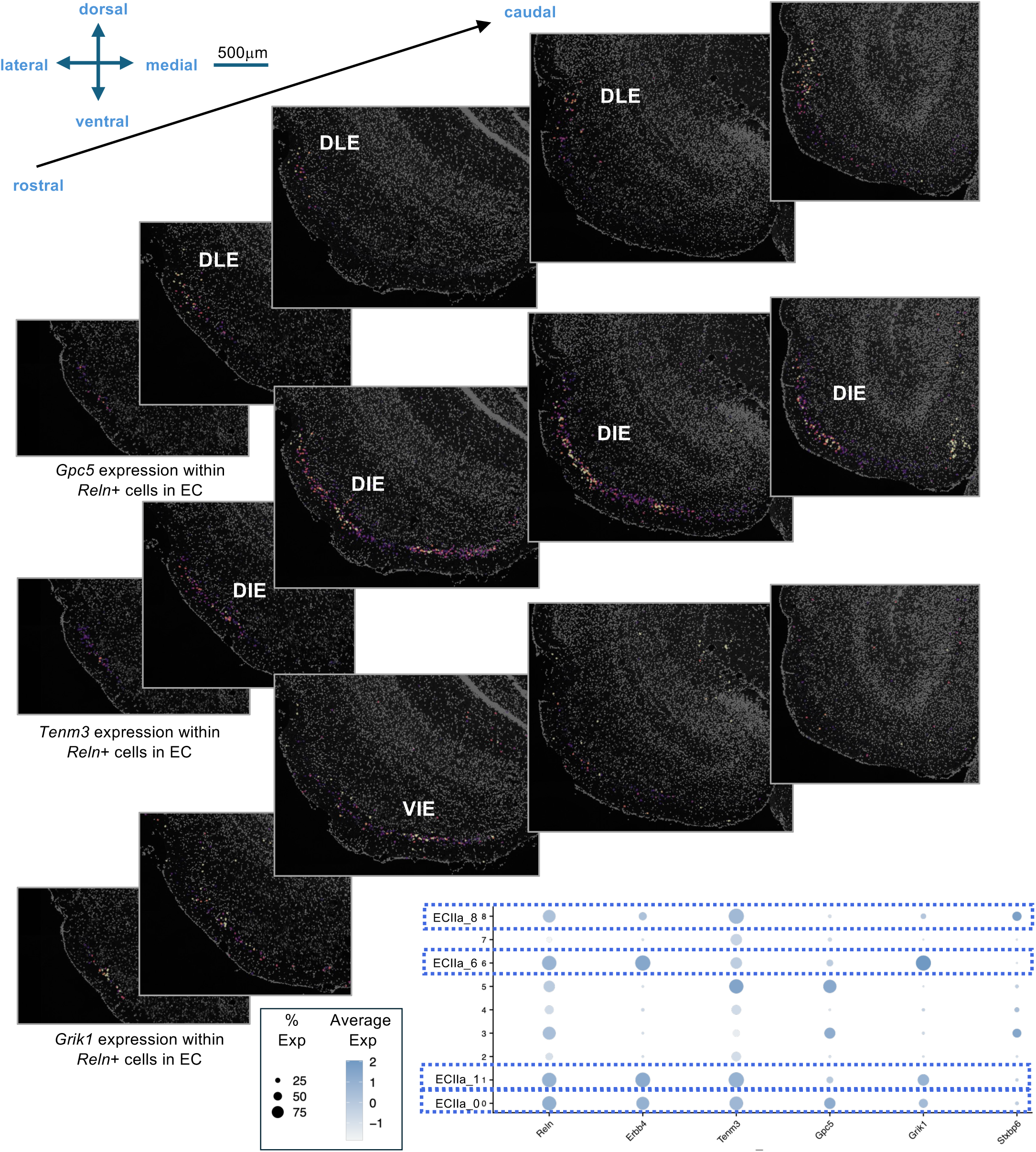
Identification of LECIIa subregions in coronal sections of a 3-month-old C57Bl/6J female mouse brain. ISH detection of *Gpc5*, *Tenm3*, *Grik1* and *Reln*. Nuclei were counterstained with DAPI. We first classified cells based on *Reln* expression. Among the classified Reln-positive cells, we then displayed heatmap of cell mean expression for *Gpc5*, *Tenm3* and *Grik1* respectively, in order to reveal the subset of ECIIa cells with highest level of expression of these genes. The Dot Plot shows expression of the four genes across ECIIa clusters. All clusters express *Reln* and *Erbb4* like all ECIIa neurons. ECIIa_0 corresponds to *Gpc5* and *Tenm3* expressing neurons an EC subarea previously referred to as DLE. ECIIa_1 neurons express *Tenm3*, an area referred to as DIE. ECIIa_6 expresses *Grik1*, referred to as VIE. ECIIa_8 expresses *Stxbp6*. The ISH reveals the position of ECIIa_0, ECIIa_1, ECIIa_6 while detection of ECIIa_8 is shown in Supplementary Figure 10. Similar results were obtained in coronal sections of a 3-month-old C57Bl/6J male mouse brain.

**Supplementary Figure 10.**
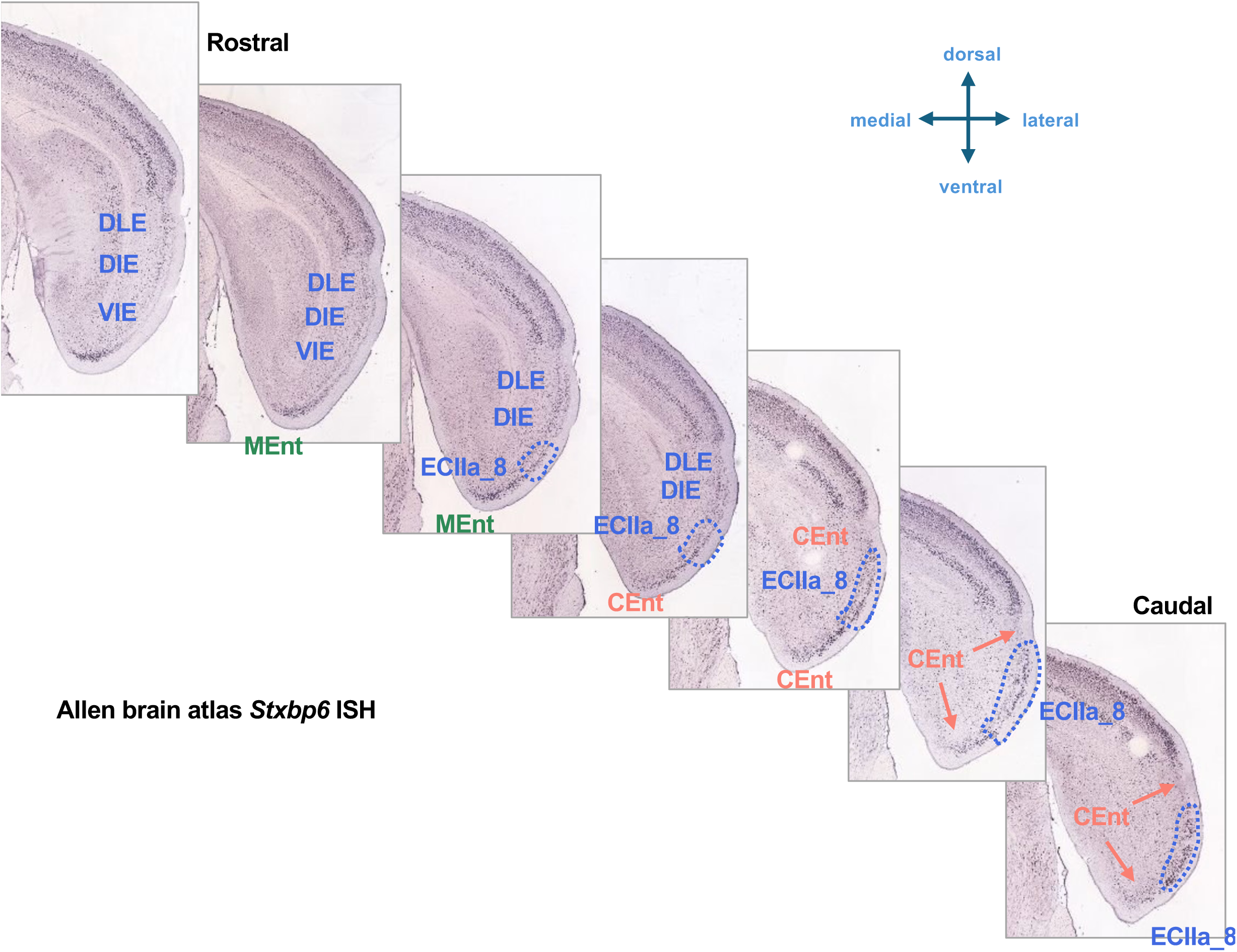
Detection of ECIIa_8. To locate LEC cluster ECIIa_8, we obtained Allen Institute ISH atlas images for *Stxbp6*, highly enriched in ECIIa_3 and ECIIa_8. The only LECIIa neurons with *Stxpb6* expression are the ones from ECIIa_8. We therefore conclude that this transition cluster is in the location outlined on these coronal sections.

**Supplementary Figure 11.**
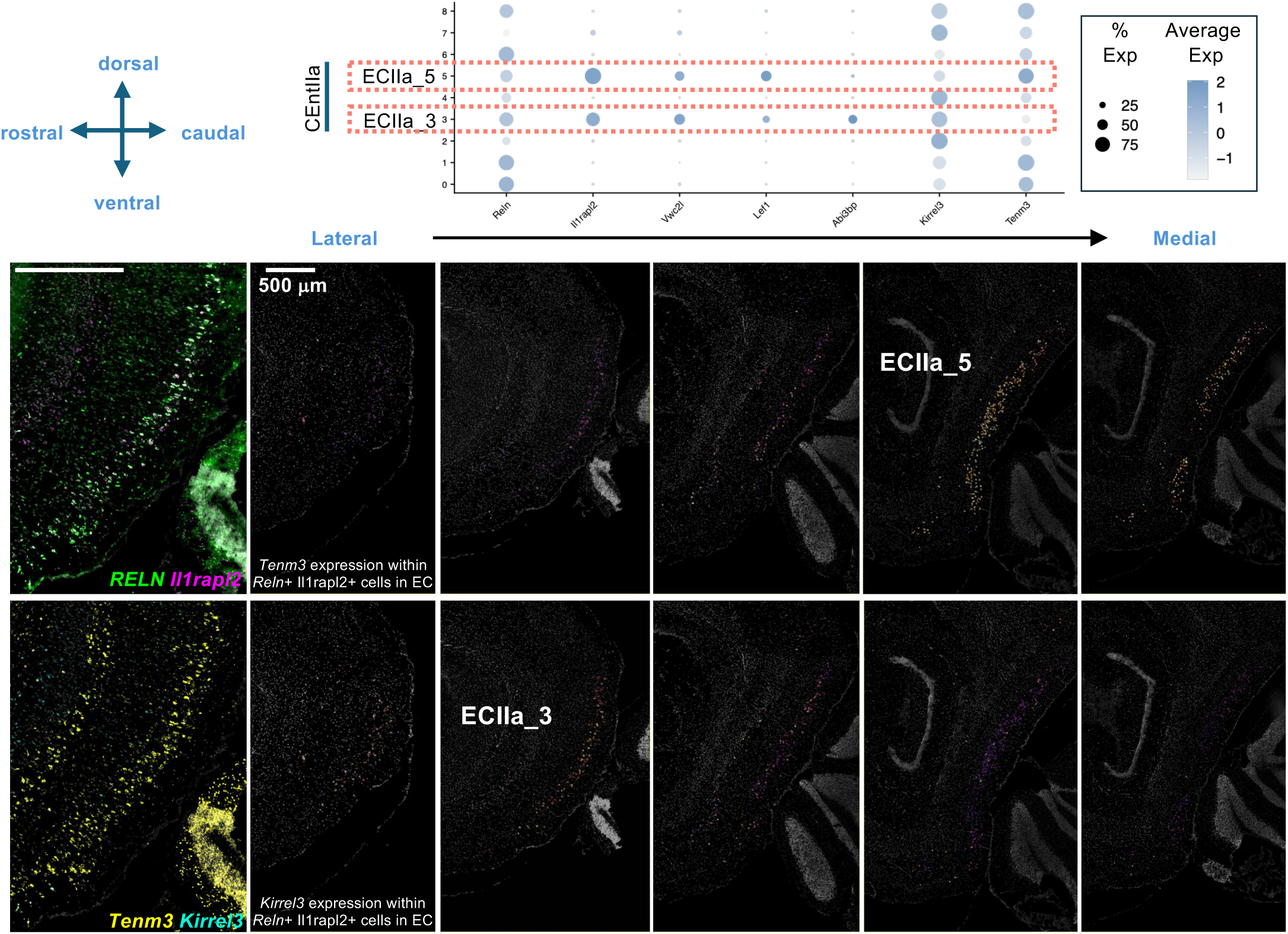
Identification of CEntIIa subregions in sagittal sections of a 3-month-old C57Bl/6J female mouse brain. Co-detection of *Kirrel3*, *Tenm3*, and *Il1rapl2* by ISH and RELN by IF. Nuclei were counterstained with DAPI. We first classified cells based on RELN and *Il1rapl2* co-expression, i.e. CEntIIa. Among the classified CEntIIa cells, we then displayed a heatmap of cell mean expression for *Tenm3* and *Kirrel3* respectively. The raw ISH signal is presented on the left column. The next five columns are a series of sagittal sections arranged from lateral-most to medial-most. The CEntIIa subarea with highest *Kirrel3* expression, ECII_3 is found on the lateral side of CEntIIa. The subarea with higest *Tenm3* expression, ECIIa_5 is found on the medial side of CEntIIa.

**Supplementary Figure 12.**
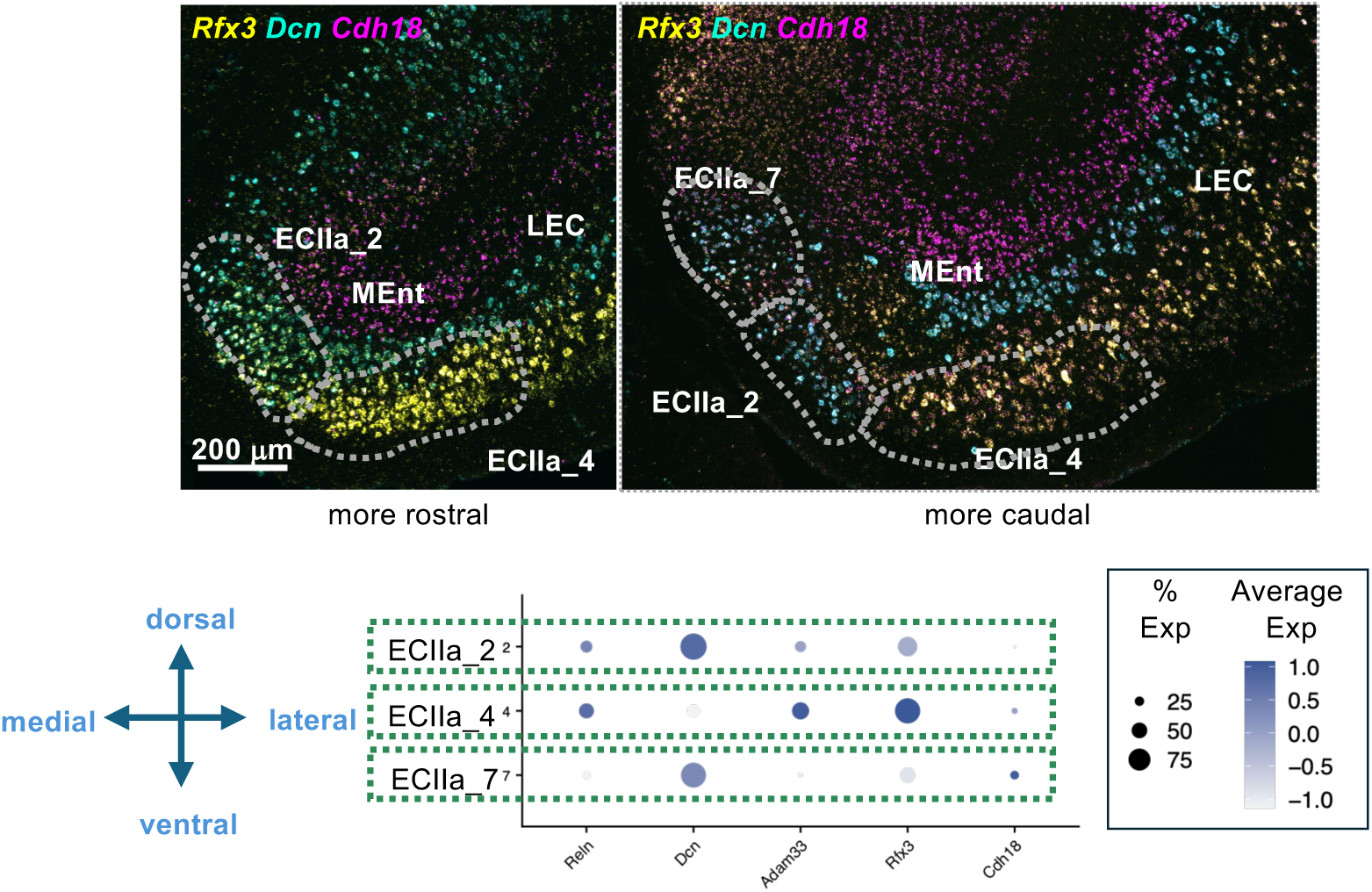
Identification of MEntIIa subregions in coronal sections of a 3-month-old C57Bl6/J female mouse brain. Close-up image of the ISH series from **Supplementary Figure 7.** ISH detection of *Rfx3* (yellow), *Cdh18* (magenta) and *Dcn* (cyan) in ISH – 3^rd^ and 4^th^ sections from the series (the 3^rd^ one is the most rostral one). The raw ISH signal is shown. The MEntIIa subarea with highest *Rfx3* expression, ECIIa_4, is the most lateral one. Both ECIIa_2 and ECIIa_7 have *Dcn* expression, found more medially. But ECIIa_7 also has expression of *Cdh18*, which we found only on the medial most subarea of MEnt.

**Supplementary Figure 13.**
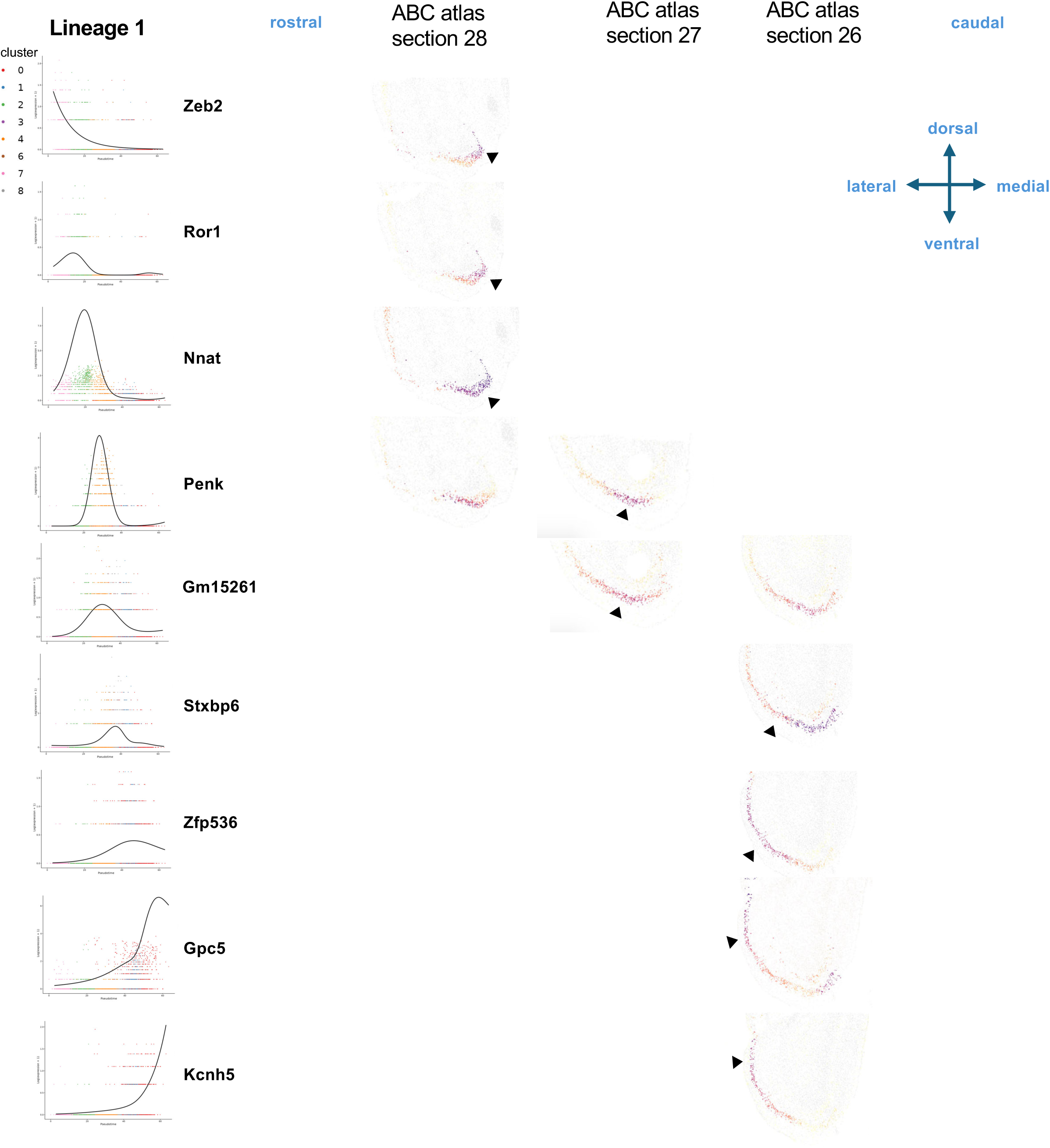
Trajectory inference analysis lineage 1. We identified genes with peak expression at various points of the lineage 1 pseudotime, and obtained the expression heatmap for these genes on coronal sections from the ABC atlas at various rostrocaudal levels where expression of the corresponding gene is maximal. We pointed with arrowheads to the anatomical position where the gene appeared to be expressed at highest level. Our data indicate that lineage 1 corresponds to an anatomical gradient that starts from the most medial parts of MEntIIa, moves laterally towards the LEC, traverses the transition cluster between MEnt and LEC (ECIIa_8), followed by DIE and finishes in DLE.

**Supplementary Figure 14.**
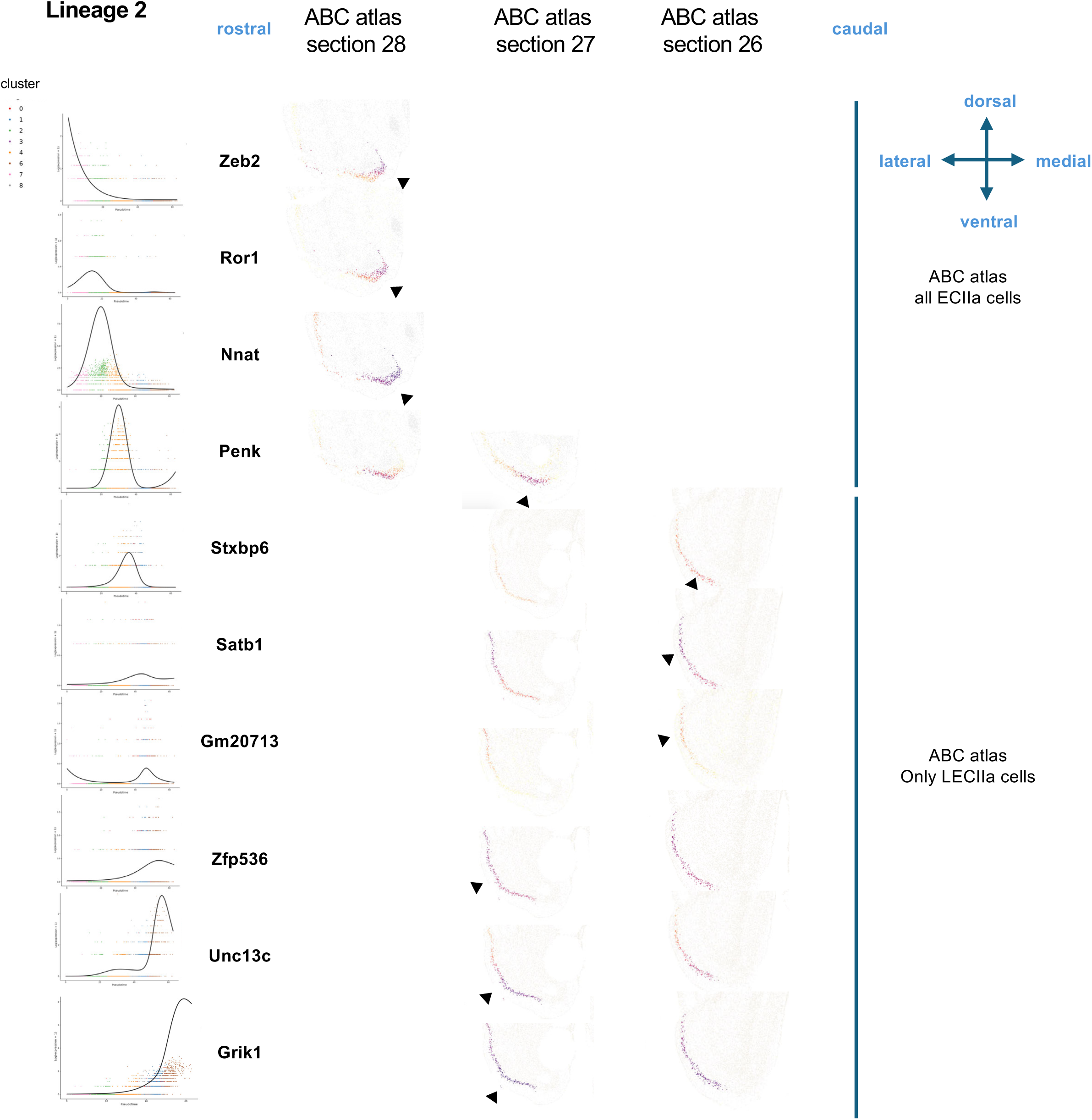
Trajectory inference analysis lineage 2. We identified genes with peak expression at various points of the lineage 2 pseudotime, and obtained the expression heatmap for these genes on coronal sections from the ABC atlas at various rostrocaudal levels where expression of the corresponding gene is maximal. We pointed with arrowheads to the anatomical position where the gene appeared to be expressed at highest level. Our data indicate that lineage 2 corresponds to an anatomical gradient that starts from the most medial parts of MEntIIa, moves laterally towards the LEC, traverses the transition cluster between MEnt and LEC (ECIIa_8), followed by DIE and finishes in VIE.

**Supplementary Figure 15.**
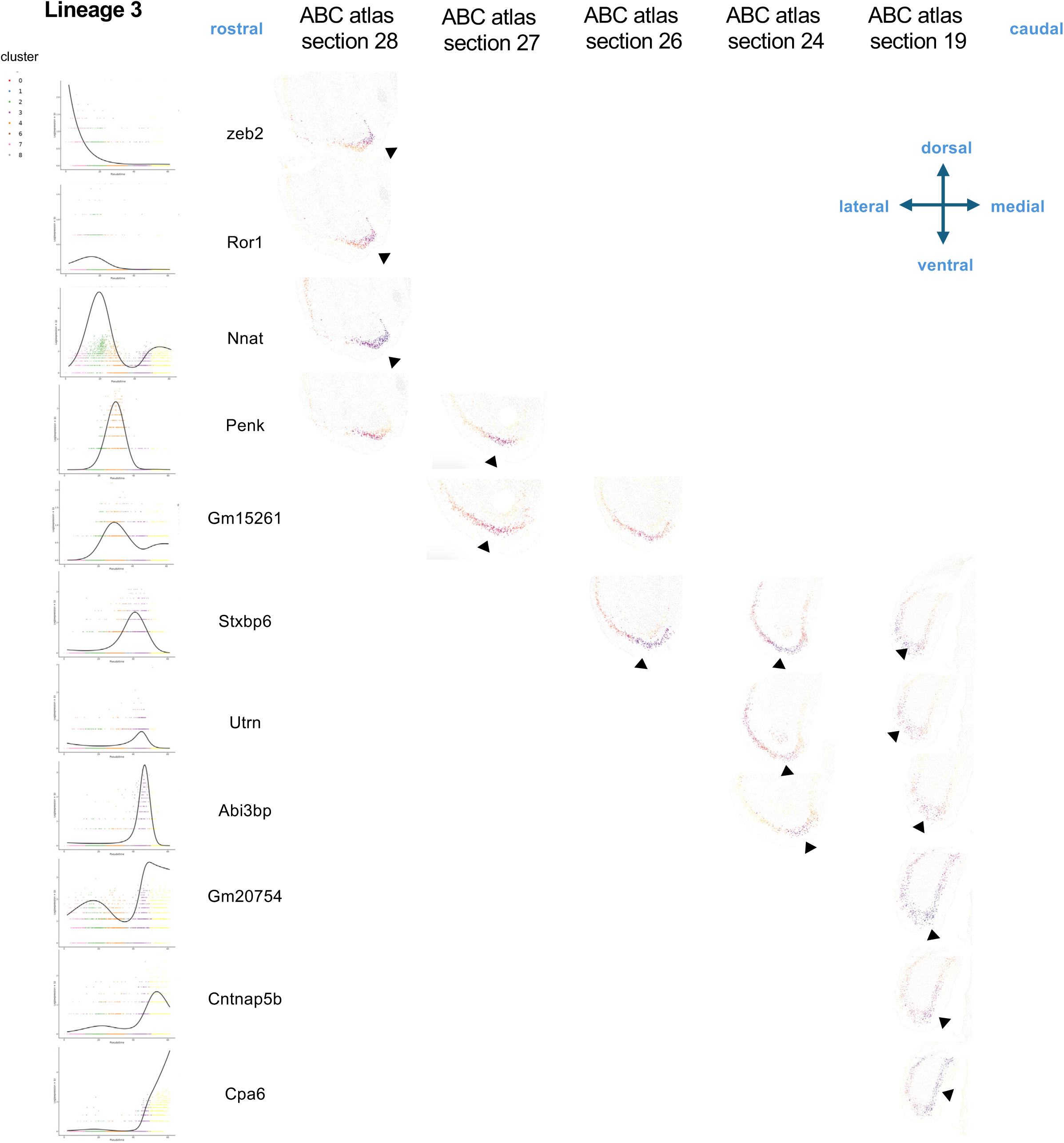
Trajectory inference analysis lineage 3. We identified genes with peak expression at various points of the lineage 3 pseudotime, and obtained the expression heatmap for these genes on coronal sections from the ABC atlas at various rostrocaudal levels where expression of the corresponding gene is maximal. We pointed with arrowheads to the anatomical position where the gene appeared to be expressed at highest level. Our data indicate that lineage 3 corresponds to an anatomical gradient that starts from the most medial parts of MEntIIa, moves laterally towards the LEC, traverses the transition cluster between MEnt and LEC (ECIIa_8), and finishes in CEnt.

**Supplementary Figure 16.**
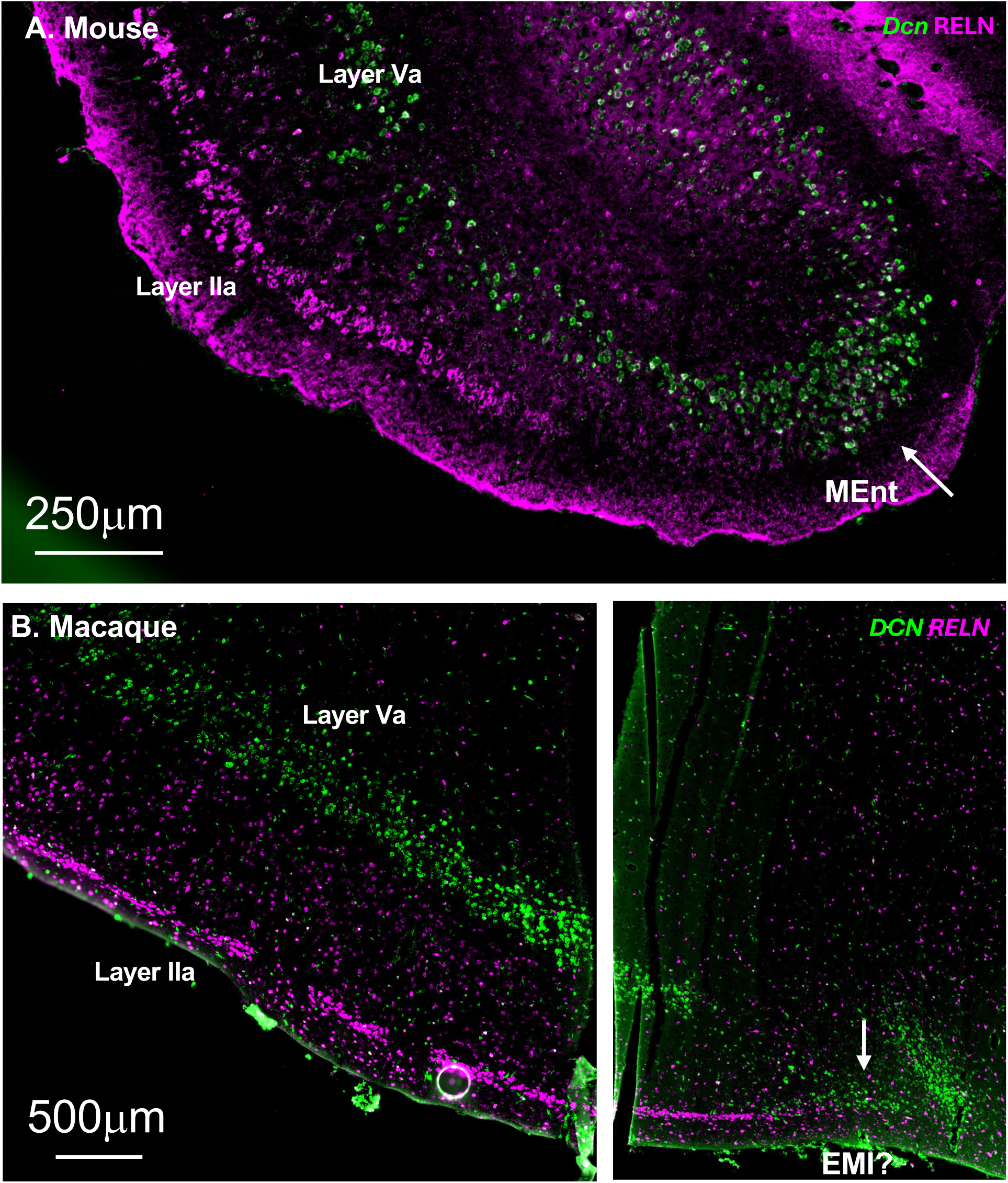
Mouse-macaque homology of MEnt. A. In the mouse, we had found that *Dcn* labels a subset of neurons from MEntIIa (Supplementary Fig. 6), labeling cells there that express only low levels of RELN. In LEC, *Dcn* labels ECVa neurons, and this layer appears to oblique towards superficial layers at the border between LEC and MEnt. In macaque, we observe a similar situation, where on the medial edge of EC, *DCN*, which is expressed in ECVa in the EC, obliques towards superficial layers. In layer II, a population is stained with *DCN* but not with *RELN*, which is not expressed anymore in layer II at this level (the section was inadvertently cut and was reconstituted on 13B). This population could represent the EMI region previously described in human.

## Supplementary Table description

**Supplementary Table 1: Cluster annotation and marker genes for all clusters in the initial cell clustering.**

**Supplementary Table 2: Cluster annotation and marker genes for all clusters in the inhibitory neuron subclustering.**

**Supplementary Table 3: Cluster annotation and marker genes for all clusters from the excitatory neuron subclustering.**

**Supplementary Table 4: Cluster annotation and marker genes for all clusters from the final_exc14 cluster subclustering.**

**Supplementary Table 5: MapMyCells output for all cells from our dataset and percentage of supertype labels identified for each of the final_exc subclusters.**

**Supplementary Table 6: GSEA pathway analysis for our comparison between ECIIa neurons and all other excitatory neurons from EC and for our pairwise comparisons LECIIa vs.**

**MEntIIa, CEntIIa vs. MEntIIa, LECIIa vs. CEntIIa.**

## Data Availability

The raw and processed sequence files generated in this manuscript are available at the gene expression omnibus (GEO) under accession number GSE349381.

## Conflict of Interest

The authors declare that the research was conducted in the absence of any commercial or financial relationships that could be construed as a potential conflict of interest.

## Author Contributions

AMM, HK, JS, SJ, DA, JPR performed the experiments. BSD analyzed all genomics data. AMM and JPR analyzed all other data. DR and CD provided tissue for the experiments. JL and JPR obtained funding. JL, DR, CD provided expert advice on the study. AMM, BSD and JPR wrote the manuscript.

## Acknowledgments

This study was supported by Cure Alzheimer’s Fund, NIA R01AG085444, R21AG085464 and R21AG084041. AMM was supported by La Caixa foundation. We thank the Rockefeller University Genomics resource center for their help with snRNAseq, Nicholas Crossland, Hans Gertje, Aoife O’Connell for their assistance with the PhenoImager, funded by NIH Award Number S10OD030269, Kathleen Rockland and Patricia Rodriguez-Rodriguez for insight on the data and manuscript.

## Notes

### Competing Interest Statement

The authors have declared no competing interest.

